# When clades collide: Genomic admixture in blacklegged ticks (*Ixodes scapularis*) from the Great Plains

**DOI:** 10.64898/2026.06.01.729354

**Authors:** Kaylee S. Herzog, Zach Pella, Halie Smith, Dayna McCormick, Lily Nuss, Amanda Brinkworth, Sujata Chaudhari, Kathryn Duncan, Kellee Sundstrom, Sydney Stein, Holly Black, Jose E. Pietri, Ryan C. Smith, Jeff Hamik, Roberto Cortiñas, Joseph R. Fauver

**Author notes:** Correspoinding author. Authors contributed equally (Kaylee S. Herzog, Zach Pella).

## Abstract

*Ixodes scapularis* ticks transmit a number of pathogens important to human health, including *Borrelia burgdorferi*, the causative agent of Lyme disease. While *I. scapularis* is found across the Eastern United States, Lyme disease transmission is largely limited to the Northeast and Upper Midwest and is nearly absent in the South. This indicates that differences in northern versus southern clades of *I. scapularis* are associated with differences in Lyme disease transmission risk. *I. scapularis* is undergoing range expansion, including into the Great Plains region of the United States. Determining where *I. scapularis* populations in the Great Plains originated can inform future risk of Lyme disease transmission in this region. In this study, we use a population genomics framework to characterize diversity, structure, and *B. burgdorferi* infection rates of *I. scapularis* populations in the Great Plains region. We generated whole genome sequence data and single nucleotide polymorphism (SNP) datasets from *I. scapularis* ticks collected in Iowa, Kansas, Nebraska, and South Dakota to compare to publicly available data from across the species’ range. Our analysis of 200 *I. scapularis* SNP datasets indicated geographically well-defined populations that correspond to historical northern and southern ancestral clades of *I. scapularis*. Ticks from South Dakota, Iowa, and northeastern Nebraska are genetically similar populations from the Upper Midwest, while ticks from Kansas are more genetically similar to ticks from the Southeast. Ticks collected in the central eastern region of Nebraska, however, represent an evenly admixed population of both northern and southern genomic backgrounds. Analysis of *B. burgdorferi* reads from genomic datasets shows ∼50% infection rate in ticks from Iowa and northeastern Nebraska, whereas ticks from Kansas show no evidence of *B. burgdorferi* infection. Of note, evenly admixed ticks from the central eastern region of Nebraska also show no evidence of *B. burgdorferi* infection. These results provide further evidence that tick genomics may influence traits associated with *B. burgdorferi* infection status, and thus the potential for Lyme disease transmission. As the range of *I. scapularis* continues to expand, bringing historically isolated populations into contact, there is a clear need to understand the consequence of genomic admixture for *B. burgdorferi* transmission potential to inform future risk of Lyme disease in the United States.

## Introduction

*Ixodes scapularis* (blacklegged tick, deer tick) is an important vector of multiple pathogens relevant to human health, including *Anaplasma phagocytophilum, Babesia microti, Ehrlichia eauclairensis*, Powassan virus, and the causative agents of Lyme disease, *Borellia miyamotoi*, and *B. burgdorferi sensus stricto* and *B. mayonii^1^*. Lyme disease is the most prevalent vector-borne disease (VBD) in the United States, and the annual number of diagnosed cases has been increasing for decades^2,3^. Unsurprisingly, the economic burden of Lyme disease in the United States is projected to be hundreds of millions of dollars annually^4^. As with most VBDs, the reported case counts are likely substantial under estimates of total infections^5^, thus making Lyme disease one of the most consequential infectious diseases in the United States. Lyme disease transmission is prevalent in the Northeast and Upper Midwest, but is nearly absent in the South^6^, highlighting important geographical differences in risk.

The tick vector of Lyme disease, *I. scapularis*, is now distributed across much of the Eastern United States, but its demographic history is characterized by dramatic range expansion and contraction^1,7^. Following deforestation and collapse of white-tailed deer populations (*Odocoileus virginianus*) across much of the Northern United States in the late 19^th^ century, *I. scapularis* populations contracted to few refugia sites in the Northeast and Upper Midwest, while populations in the Southeast persisted. Following reforestation and recovery of deer populations, these refugia seeded contemporary northern populations of *I. scapularis*^*7*,*8*^. Dramatic expansion of these ancestral populations has since occurred in the Northeast and Upper Midwest, as well as in the Southeastern United States since the 1970s. However, the increased burden of Lyme disease has largely been limited to the Northeast and Upper Midwest^9,10^. Geographic differences in Lyme disease epidemiology can be attributed, in part, to regional differences in tick populations. Multiple studies have demonstrated *I. scapularis* populations have undergone isolation by distance and that gene flow is regionally restricted, resulting in genetically distinct northern and southern clades^8,11–15^. Life history traits in nymphal stage ticks important for *B. burgdorferi* transmission, such as questing behavior and host preference, also differ by geography. Ticks collected in the Northeast and Upper Midwest quest above the leaf litter and prefer to feed on small mammals that maintain *B. burgdorferi* transmission cycles, whereas *I. scapularis* populations from the Southeast quest below the leaf litter and preferentially feed on reptiles, which are incompetent hosts for *B. burgdorferi^16–22^.* These life history traits result in differences in nymphal infection rates and ultimately, differences in human Lyme transmission risk^9^. Thus, contemporary *I. scapularis* populations collected in the Northeast and Upper Midwest are genetically and behaviorally distinct from ticks collected in the Southeast.

The range of *I. scapularis* is expanding into previously uninhabited regions, including the Great Plains. While populations of *I. scapularis* have been known for decades from Iowa (1989)^23^ and Kansas (1988)^24^, established populations have recently been identified in North Dakota (2010)^25^, South Dakota (2015)^26^, and Nebraska (2018)^27^, representing the western-most edge of the species’ distribution^7^. Additionally, *B. burgdorferi* has been detected in *I. scapularis* ticks from all three of these states^25,28–30^. Following the 2021 identification of autochthonous Lyme disease transmission in Nebraska, expanded tick surveillance identified two foci in northeastern and central eastern regions of the state. Molecular surveillance for *B. burgdorferi* in Nebraska demonstrates that *I. scapularis* from the northeastern region have a 55.4% prevalence in questing ticks (N=119 tested), whereas ticks in the central eastern foci have a prevalence of 0% (N=113 tested)^31^. Clear differences in *B. burgdorferi* prevalence in questing ticks in Nebraska led us to test the hypothesis that the geographic structure of populations observed for *I. scapularis* in the Eastern United States is also present in the Great Plains and that Nebraska is a potential contact zone between historically isolated northern and southern populations of *I. scapularis*.

In this study, we used whole genome sequencing (WGS) to characterize the genomic similarity, population structure, and connectivity within and among recently-established populations of *I. scapularis* in the Great Plains. We compared these newly-generated datasets to WGS data from *I. scapularis* across the Eastern United States to better understand the genetic background of recently-established populations. We also used WGS to explore the presence or absence of *B. burgdorferi* in individual ticks. Understanding the genetic ancestry of established ticks, how populations are structured, and what phenotypic differences exist between them can inform the risk of Lyme disease transmission in the Great Plains and other areas of recent *I. scapularis* range expansion.

## Methods

### Specimen collection and preservation

Sampling locations for ticks collected in this study are presented as open circles in **Figure 1**, and locations of sampling from publicly available datasets are shown in closed circles. Adult *I. scapularis* ticks from recently-established populations in South Dakota and Nebraska were prioritized for WGS. Additionally, samples collected from more established populations in Iowa and Kansas were sequenced to provide contextual data from populations of *I. scapularis* in close proximity and to add geographic contiguity among states previously sampled across the region. Detailed metadata for all specimens included in analyses, including sampling locality, life-cycle stage, nucleic acid isolation methodology, sequencing and alignment summary statistics, etc. are presented in **Supplemental Table 1**. Host seeking specimens were collected in Nebraska (n=37) using standardized drag sampling during fall 2024 and spring 2025 in eastern Nebraska. Dragging was conducted for approximately 1–4 hours per site (mean ∼2 hours), and drag cloths were examined for ticks at 25 m intervals. All collected ticks were preserved in 70% ethanol and identified to species and life stage in the laboratory. Additionally, deer checks were conducted during fall 2024 at designated check stations to assess host-associated tick presence. All ticks collected from a single deer were placed into a vial containing 70% ethanol, and tick species, sex, life stage, and total counts were recorded per deer. All infested deer were killed in Dodge, Washington, and Douglas counties. Host seeking specimens from Iowa (n=19) were collected as described for Nebraska using active tick drags in April or June of 2023. Live ticks were placed in vials and stored at -80°C prior to shipping to UNMC. Host seeking ticks were collected in South Dakota (n=5) by flagging vegetation. Following species identification, ticks were cut in half with a scalpel and frozen at -20°C or -80°C prior to shipping to UNMC. For specimens from Kansas (n=5), field collection and preservation methods were previously described^32,33^. In addition to field-collected specimens, live nymphal ticks (n=3) were obtained through BEI Resources from a laboratory strain originally established using ticks collected from Rhode Island. Nymphs were killed via freezing at -80°C prior to homogenization and gDNA purification.

**Figure 1.**
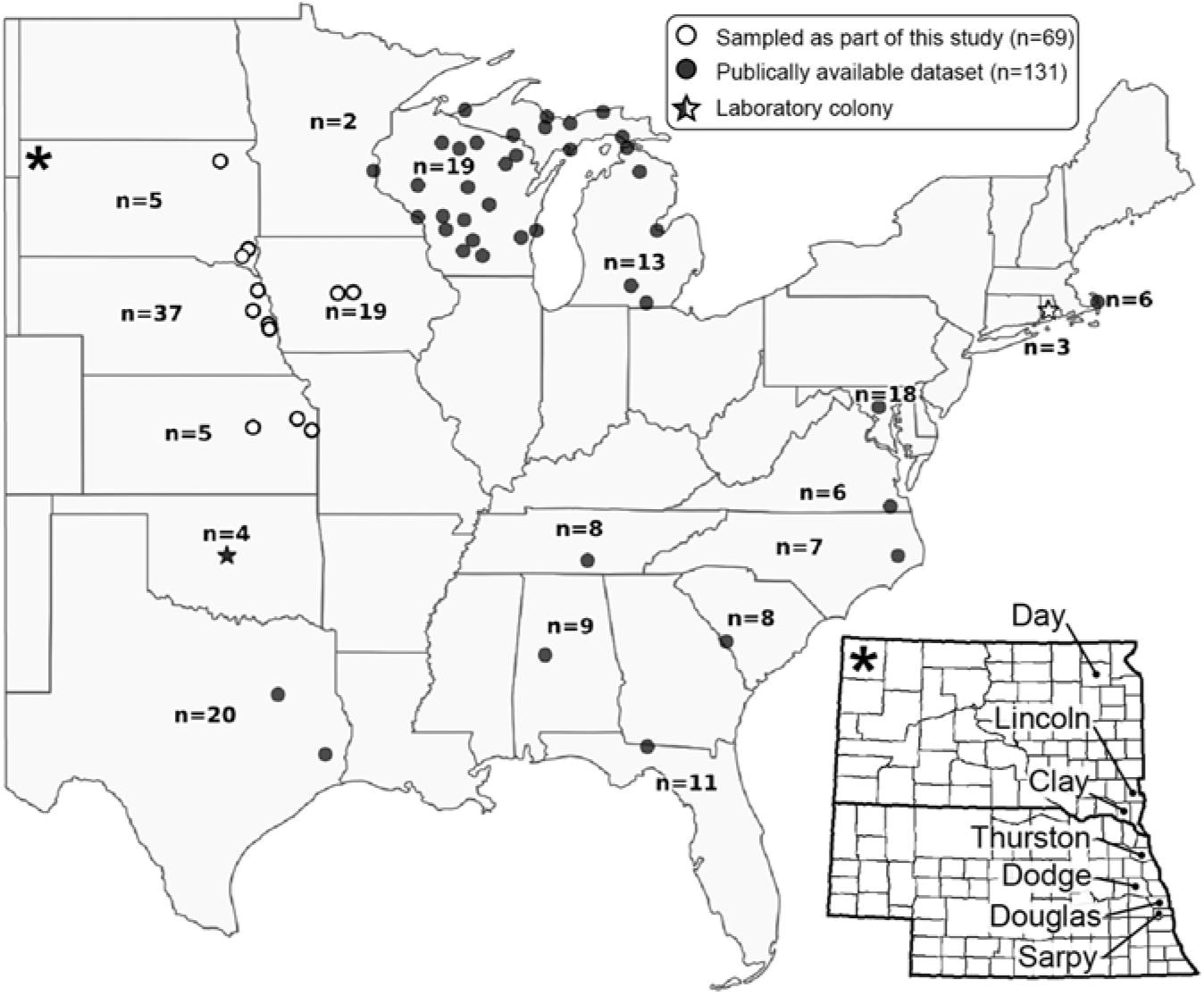
Sampling localities for individuals of *Ixodes scapularis* included in population genomic analyses. Circles indicate field collection sites; stars indicate laboratory colonies. Open shapes indicate localities that were newly-sampled as part of this study; closed shapes indicate localities for publically-available datasets that were downloaded for inclusion. Values of n indicate the total number of individuals of *I. scapularis* included from each state. Detailed map at right (denoted by asterisk) shows county-level designations for each sampling site in the Great Plains (South Dakota and Nebraska).

### Sample processing

Individual ticks underwent surface sterilization using sequential washes of 3% H_2_O_2_ (1 minute), 70% ethanol (two 30-second washes), and 1× PBS (2 minutes) with vortexing between each step. Sterilized ticks were homogenized in 100μL phosphate buffered saline (PBS; 1X; pH 7.4; Thermo Fisher Scientific) in Safe-Lock Tubes (Eppendorf) using a TissueLyser II tissue disruptor (QIAGEN) with stainless steel beads at 25 Hz for 5, 10, or 30 min (see **Supplemental Table 1**). Total genomic DNA was purified from homogenized tissue with two approaches. The first approach used the DNeasy Blood & Tissue Kit (Qiagen) following the manufacturer’s protocol with overnight proteinase K digestion at 56°C and elution in 50µL AE Buffer. The second approach used the Zymo Quick DNA HMW MagBead Kit (Zymo Research) and the modified purification protocol previously described^34^. For ticks extracted using the Zymo Kit, homogenization was done in 95 µL Zymo Biofluid & Solid Tissue Buffer rather than PBS. Purified gDNA from both approaches was quantified using a Qubit™ double-stranded high sensitivity DNA assay (dsDNA HS; Invitrogen). For the specimens from Kansas (n=5; see **Supplemental Table 1**) gDNA was previously extracted as described by McClung et al (2023)^33^. Prior to library preparation, these extractions were bead-cleaned using KAPA Pure Beads (Roche) in a 1:1 ratio, eluted in nuclease-free water, and quantified using Qubit™ dsDNA HS assay.

### Next-generation sequencing library preparation

For each specimen, 15–40 ng of gDNA was used as input to library preparation (see **Supplemental Table 1**). Individual WGS libraries were prepared using an Illumina DNA Prep Kit (Illumina) following the manufacturer’s protocol with 6 or 8 cycles of unique dual-indexing PCR, depending on gDNA input (see **Supplemental Table 1**). Purified indexed libraries were quantified via Qubit™ dsDNA HS assay and pooled for equal concentration. Pools then underwent an additional round of double-sided bead purification following the ratios and incubation times given in the Illumina DNA Prep Kit protocol. Bead-purified pools were quantified via Qubit™ dsDNA HS assay and the KAPA Library Quantification Kit for Illumina Platforms (KAPA Biosystems). Insert length was assessed with High Sensitivity D1000 DNA ScreenTape analysis (Agilent) on an Agilent 2200 TapeStation system.

### Next-generation sequencing

Pooled libraries were submitted to the University of Nebraska Medical Center Genomics Core Facility (UNMC GCF) for 2×150 paired-end Illumina sequencing in two rounds. For the first round, the bead-purified pool (n=61 libraries) was sequenced on an Illumina NovaSeq 6000 S2 flow cell. For the second round, the bead-purified pool (n=8 libraries) was sequenced on an Illumina NovaSeq 6000 S1 flow cell. For each round, 10X depth of coverage (DOC) across the *I. scapularis* genome was targeted per individual. Data were demultiplexed by the UNMC GCF and raw reads from the same libraries sequenced across multiple lanes were concatenated prior to analysis.

### Integration of previously generated data

To establish a comprehensive population framework and contextualize our newly generated data, we integrated publicly available WGS Illumina data from the National Center for Biotechnology Information Sequence Read Archive (NCBI SRA). For *I. scapularis*, 131 datasets were selected, providing coverage across 13 states representing the established northern and southern clades (**see Figure 1, Supplemental Table 1**). Raw reads were downloaded using fasterq-dump through the NCBI SRA Toolkit v2.11^35^.

### Data quality control and reference alignment

Illumina sequence datasets generated herein (n=69) and downloaded from the NCBI SRA (n=131) were processed using the same bioinformatic pipeline (see **Supplemental Figure 1**). Raw reads were quality-filtered using fastp v0.23^36^ to trim adaptor sequences and remove low-quality reads and reads shorter than 50 base pairs (bp). FastQC v0.12^37^ was then used to confirm successful trimming and filtration. Repeat regions in the *I. scapularis* reference genome (ASM1692078v2)^38^ were masked using RepeatMasker v4.0^39^ and then quality-controlled reads were aligned to the masked reference using BWA-MEM v0.7^40^ with default parameters. To minimize potential biases arising from uneven sequencing depth across samples, the eight libraries sequenced on an S1 flow cell that achieved substantially higher coverage (∼30X) than the target depth were downsampled to match the coverage distribution of the remaining samples (∼8X). Downsampling fractions were calculated for each sample based on original read counts, targeting approximately 135 million reads per sample. Downsampling was performed using samtools v1.20 with the -s flag, which randomly subsamples reads using a fixed seed (seed=42) to ensure reproducibility. Resulting sequence alignment map (SAM) files were converted to BAM format, coordinate-sorted, and indexed using samtools v1.19^41^. Read groups were then added using Picard v3.0^42^ AddOrReplaceReadGroups and PCR duplicates were removed using Picard MarkDuplicates with the “--OPTICAL_DUPLICATE_PIXEL_DISTANCE 2500” flag appropriate for patterned flow cells.

### Variant detection and single nucleotide polymorphism isolation and filtration

Variant calling was performed using the Genome Analysis Toolkit (GATK) v4.6^43^ following the best practices workflow for germline short variant discovery^44,45^. First, HaplotypeCaller in GVCF mode with diploid genome modeling was used to generate gVCFs for individual samples. This process was run across scattered genomic intervals using SplitIntervals with a scatter count of 20. Resulting gVCFs were consolidated into a GenomicsDB workspace using GenomicsDBImport. Joint genotyping across all samples was performed using GenotypeGVCFs. Scattered VCFs from all samples were then gathered into a final jointly-called variant dataset using GATK GatherVcfs. Indels were removed using SelectVariants to retain only single nucleotide polymorphisms (SNPs) for downstream analysis. Baseline quality filters were applied following GATK best practices, flagging variants that failed any of the following criteria: QD < 2.0, QUAL < 30.0, SOR > 3.0, FS > 60.0, MQ < 40.0, MQRankSum < -12.5, or ReadPosRankSum < -8.0. Flagged variants were then removed using SelectVariants. Additional site-level filtration was performed using VCFtools v0.1.16^46^ to retain only variants with MAF >0.1, MAC 2, biallelism, site-level missingness >5%. Filtered datasets were then used for downstream analysis with standard population genomic software, some of which required additional filtration (see **Supplemental Figure 1**, and below).

### Population summary statistics

Summary statistics and corrected F_ST_ values were generated using the “populations” module in STACKS v2.68^47,48^. The module was run using a population grouping schemes where each state was considered its own population, with Nebraska specimens designated as either a northeastern subpopulation or a central eastern subpopulation (n=19 subpopulations total; see **Supplemental Table 1**). At least 80% of specimens within a subpopulation were required to have a genotype call in order to process a locus for that subpopulation, and at least 13 of 19 subpopulations were required to be represented at a locus for inclusion. Kernel smoothing was enabled for F-statistic calculation and F_ST_ correction was implemented with a p-value cutoff of 0.05.

### Principal components-based analyses

Principal components analysis (PCA) was conducted using PLINK v2.0a1^49^. For PCA, the filtered SNP dataset was converted to binary format and linkage disequilibrium (LD) pruning was performed using a sliding window approach (window size of 50kb, window step size of 10bp, and r^2^ threshold of 0.1). The resulting PCA plot was visualized using matplotlib v3.10.0^50^ and seaborn v0.13.2^51^. Discriminant analysis of principal components (DAPC) was performed using the R package adegenet v2.1.11^52,53^ in R v4.5.1^54^. For DAPC, the most likely number of populations was determined using both Bayesian information criterion (BIC) and Akaike information criterion (AIC) and the number of retained principal components was determined through a-score optimization.

### Admixture analyses

Admixture analysis was performed using fastSTRUCTURE v1.0^55^ ADMIXTURE v1.3^56^. To ensure that fastSTRUCTURE could be reliably used in place of STRUCTURE v2.3^57,58^, a preliminary analysis was performed on a reduced dataset to compare outputs from both programs. For both reduced and full datasets, VCFs were first filtered for LD as described above using PLINK2, then converted to STRUCTURE format using PGDSpider v2.1.1.5^59^ with a custom SPID configuration file specifying diploid genotype handling, SNP-only data export, and exclusion of monomorphic SNPs. Missing data were preserved and coded as “-9”. No population definition file was used during conversion to allow for *de novo* inference of admixture. Following conversion, the STRUCTURE format file was manually edited to remove the first row containing SNP identifiers as these labels can interfere with proper file parsing. For STRUCTURE using the reduced dataset, k-values 1–5 were tested with 10 independent runs completed for each k-value, with 250,000 generations for each run with the first 50,000 generations discarded as burn-in. The most likely k-value was determined using likelihood scores and the Evanno ΔK method^60^ implemented in STRUCTURE HARVESTER v.0.6.93^61^. For fastSTRUCTURE using the reduced and full dataset, k-values 1–10 were tested using the simple prior with default convergence criterion and the full likelihood model. The optimal k-value was determined using two complementary approaches implemented in the chooseK.py script: (1) the k-value that maximized the variational lower bound (marginal likelihood), and (2) the modal k-value across runs based on the number of model components required to explain structure in N-1 individuals from the admixture proportions (Q matrices). Population membership coefficients were visualized using the district.py script included with fastSTRUCUTRE, with samples grouped and ordered by state. For ADMIXTURE, only the full LD-pruned PLINK2 fileset (.bed/.bim/.fam) was used as input; however, because ADMIXTURE requires integer chromosome codes, scaffold names were converted to sequential integers prior to analysis using a custom mapping file. ADMIXTURE was run for k=1–6 using 8 threads, with 5-fold cross-validation (--cv) enabled to assess model fit. The optimal k-value was determined by identifying the k with the lowest cross-validation error. Ancestry proportion estimates (Q matrices) were visualized with samples grouped by state for comparison with fastSTRUCTURE results.

### Isolation by distance visualization

To visualize isolation by distance (IBD), the geographic distance between populations was calculated using the Haversine formula and plotted against pairwise corrected F_ST_ values generated via STACKS (see above). For populations represented by two or three sampling localities (Iowa, Kansas, central eastern Nebraska, South Dakota, Texas), the geographic coordinates of the midway point among sampling localities were used for distance calculations, and for populations represented by more than three sampling localities (Michigan, Wisconsin) the geographic coordinates for the centroid of the state were used. Each state-based population was characterized as either “majority northern descent” or “majority southern descent” based on results from principal component- and admixture-based analyses, with the central eastern Nebraska population defined as evenly split between the two. Pairwise comparisons representing majority northern descent versus majority southern descent populations were highlighted separately from other comparisons. Populations represented by laboratory colonies (Rhode Island–BEI, Oklahoma–OSU) were excluded from IBD estimations.

### Determination of *Borrelia burgdorferi* infection rates from whole genome sequencing data

To identify reads from associated *B. burgdorferi* bacteria, NGS were aligned to a multireference file that contained both the *I. scapularis* reference genome (ASM1692078v2) and the *B. burgdorferi* reference genome (GCA_000181575.2, Bol26). Reads were aligned using BWA-MEM v0.7 and regions corresponding to *B. burgdorferi* were extracted to .bam files and filtered with samtools view for alignment quality with following flags: -q 20 -e ‘(qlen-sclen)>75’. The number of reads aligning to each *B. burgdorferi* genome/plasmid sequence, and the total number of reads aligning to *B. burgdorferi* overall were calculated for each sample using samtools flagstat. Identifying multiple reads across multiple genomic and/or plasmid contigs was used as an indication for infection with *B. burgdorferi*.

## Results

### DNA extraction and library preparation

In total, we generated 69 novel WGS datasets from individual specimens of *I. scapularis*, collectively from Nebraska (n=37), Iowa (n=19), Kansas (n=5), South Dakota (n=5), and the BEI Resources Rhode Island-derived laboratory strain (n=3) (**Figure 1, Supplemental Table 1en individuals with inferr**). The specimens from Nebraska represent both the northeastern (Thurston County, n=11) and central eastern (Dodge, Douglas, and Sarpy Counties; n=26) regions of the state. Specimens from Nebraska, Iowa, Kansas, and South Dakota represent the first complete WGS data for *I. scapularis* generated from each of these states. DNA extraction efficiency differed between methodological approaches and life-cycle stages. Median gDNA yield from adult ticks extracted with the Qiagen DNeasy Blood & Tissue kit was 140.0ng, with substantial variation across samples. In contrast, nymphal ticks and sets of eight legs from single adult ticks extracted with the Zymo Quick-DNA HMW MagBead Kit yielded medians of 106.0ng and 70.8ng of gDNA, respectively, with lower among-sample variance from substantially smaller input tissue mass (**Supplemental Figure 2**).

### Next-generation sequencing, incorporation of publicly-available data, and variant calling

Sequencing of the 69 WGS libraries yielded 12.6 billion read pairs total, with 102.1–637.7 million read pairs generated per specimen (mean 182 million read pairs). Quality filtration with fastp resulted in a high proportion of data retention (95.8– 97.5%; 96.5% ± 0.4%), with low duplication rates (5.0–8.2%; 6.2% ± 0.7%). Following alignment to the repeat-masked *I. scapularis* reference genome, mapping rates were consistently high (90.7–99.0%; 98.5% ± 1.0%). Mean depth of coverage was 10.8× ± 7.3× (6.1–37.7×; median 8.4×) with 55.8% ± 9.4% of the genome covered at >5× and 29.8% ± 16.0% at >10×. To establish a comprehensive population framework, we integrated 131 publicly available WGS datasets from the NCBI SRA, representing 13 states across the established range of *I. scapularis* (**Figure 1**). These datasets yielded 10.6 billion raw read pairs total, with 32.4–701.6 million read pairs per specimen (mean 80.7 million read pairs). Data retention after quality filtration was high (96.0–99.9%; 99.3% ± 0.8%) though duplication rates were more variable as compared to the datasets generated as part of this study (5.8–23.7; 10.9% ± 3.5%). Mapping rates were comparable to those of the newly-generated datasets (96.1–99.3%; 98.6% ± 0.6%). Mean depth of coverage ranged from 2.0-10.5× (4.0× ± 1.4×; median 3.9×), with 21.8% ± 10.7% of the genome covered at >5× and 6.2% ± 5.7% of the genome covered at >10×. Joint variant calling across all 200 specimens resulted in 527,287,655 SNPs. Following stringent quality filtration using GATK and VCFtools, 92,326 high-quality SNPs were retained. Linkage disequilibrium pruning with PLINK2 further reduced the dataset to 20,914 SNPs for PCA- and admixture-based analyses (**Table 1**).

**Table 1.** Number of single nucleotide polymorphisms (SNPs) retained after each step of SNP filtration. For PCA, DAPC, fastSTRUCTURE and ADMIXTURE analysis, the fully-filtered dataset through linkage disequilbrium filtration was used (n=20,914 SNPs); for generation of population-level summary statistics using the STACKS populations module, no linkage disequilibrium filter was applied (n=92,326 SNPs).

| Unfiltered | GATK baseline quality filtration | MAF >0.1, MAC 2, biallelism, site-level missingness >5% | Linkage disequilibrium |
| --- | --- | --- | --- |
| 527,287,655 | 423,746,161 | 92,326 | 20,914 |

### Population-level summary statistics

Values for proportion of polymorphic sites, observed heterozygosity (H_O_), expected heterozygosity (H_E_), inbreeding coefficients (F_IS_) and nucleotide diversity (π) considering variant and fixed sites for all subpopulations are presented in **Table 2**. As expected, a greater proportion of polymorphic sites was generally identified in larger subpopulations. Exceptions include Texas and Florida, which are each relatively large subpopulations (>10 specimens per site analyzed on average) but had comparatively low proportions of polymorphic sites (<58%). Observed heterozygosity and nucleotide diversity were generally lower for subpopulations in southern geographic regions (H_O_: 0.094–0.266, π: 0.129–0.232) as compared to those in northern regions (H_O_: 0.193–0.408, π: 0.242–0.345), suggesting greater standing genetic diversity in northern regions. Ranges of inbreeding coefficients also indicate more inferred inbreeding in southern populations as compared to northern populations (-0.065–0.094 versus -0.168–0.191, respectively). It is worth noting that the most negative inbreeding coefficient among northern subpopulations (-0.168) was unsurprisingly inferred for the Rhode Island-derived BEI laboratory colony. For specimens from central eastern Nebraska relatively high values for observed heterozygosity and nucleotide diversity were inferred.

**Table 2.** Population-level summary statistics generated from *Ixodes scapularis* single nucleotide polymorphism (SNP) datasets (n=92,326 SNPs). Individuals were assigned to populations by state of origin, with individuals from Nebraska desginated as either a northeastern region (NE-NE; Thurston county) or east central region (NE-EC; Dodge, Douglas or Sarpy counties) population. Individuals from laboratory colonies (Rhode Island, BEI; and Oklahoma, OSU) are labeled with their associated laboratory strain acronyms.

|  | Population | Avg no. individ./site | No. sites | % polymorphic sites | H<sub>O</sub> | H<sub>E</sub> | F<sub>IS</sub> | π |
| --- | --- | --- | --- | --- | --- | --- | --- | --- |
| North | RI (BEI) | 3.0 | 90,010 | 57.92% | 0.391 | 0.244 | -0.168 | 0.292 |
|  | MN | 2.0 | 90,373 | 59.77% | 0.408 | 0.259 | -0.095 | 0.345 |
|  | WI | 18.2 | 90,986 | 82.67% | 0.293 | 0.281 | 0.025 | 0.288 |
|  | MI | 12.6 | 91,192 | 87.87% | 0.311 | 0.287 | 0.008 | 0.298 |
|  | MA | 5.7 | 86,025 | 70.34% | 0.257 | 0.246 | 0.044 | 0.270 |
|  | MD | 16.5 | 81,752 | 82.01% | 0.193 | 0.235 | 0.191 | 0.242 |
|  | IA | 18.8 | 92,096 | 94.83% | 0.402 | 0.319 | -0.153 | 0.328 |
| Great Plains | SD | 4.9 | 92,022 | 82.86% | 0.402 | 0.302 | -0.125 | 0.336 |
|  | NE-NE | 10.9 | 92,093 | 90.75% | 0.408 | 0.313 | -0.161 | 0.328 |
|  | NE-EC | 25.6 | 92,129 | 97.37% | 0.343 | 0.292 | -0.093 | 0.297 |
| South | KS | 4.9 | 91,966 | 60.88% | 0.266 | 0.208 | -0.065 | 0.232 |
|  | TN | 7.7 | 87,450 | 68.88% | 0.206 | 0.204 | 0.052 | 0.218 |
|  | VA | 5.8 | 90,435 | 53.01% | 0.203 | 0.175 | -0.013 | 0.191 |
|  | NC | 6.7 | 88,492 | 49.71% | 0.170 | 0.158 | 0.017 | 0.171 |
|  | SC | 7.7 | 86,347 | 51.21% | 0.170 | 0.162 | 0.029 | 0.173 |
|  | AL | 8.7 | 86,004 | 51.03% | 0.176 | 0.163 | 0.010 | 0.173 |
|  | FL | 10.5 | 90,696 | 54.39% | 0.173 | 0.163 | 0.020 | 0.171 |
|  | TX | 18.6 | 90,553 | 57.48% | 0.140 | 0.154 | 0.094 | 0.158 |
|  | OK (OSU) | 4.0 | 54,229 | 32.05% | 0.094 | 0.113 | 0.077 | 0.129 |

### Principal components-based analyses

PCA revealed clear population structure among the 200 individual *I. scapularis*. The first two principal components explained 11.2% and 7.9% of the total variance in the dataset, respectively, and differentiated samples into two primary clusters corresponding to northern and southern geographic origins (**Figure 2**). Specimens from northern states (Massachusetts, Rhode Island [BEI], Maryland, Michigan, Wisconsin, and Minnesota) grouped together, as did specimens from southern states (Texas, Oklahoma [OSU], Alabama, Virginia, North Carolina, South Carolina, and Florida). Specimens from Tennessee showed intermediate grouping between the two clusters, though with closer identity to the southern cluster. Newly-sequenced specimens (depicted with triangles in Figure 2) showed varying placement. Specimens from South Dakota and Iowa clustered with the northern clade while specimens from Kansas clustered with the southern clade. Among specimens from Nebraska, geographic sub-structuring was evident. Ticks from northeastern Nebraska (Thurston County) grouped with the northern cluster, while ticks from central eastern Nebraska (Dodge, Douglas, and Sarpy Counties) showed intermediate positioning between the two clusters.

**Figure 2.**
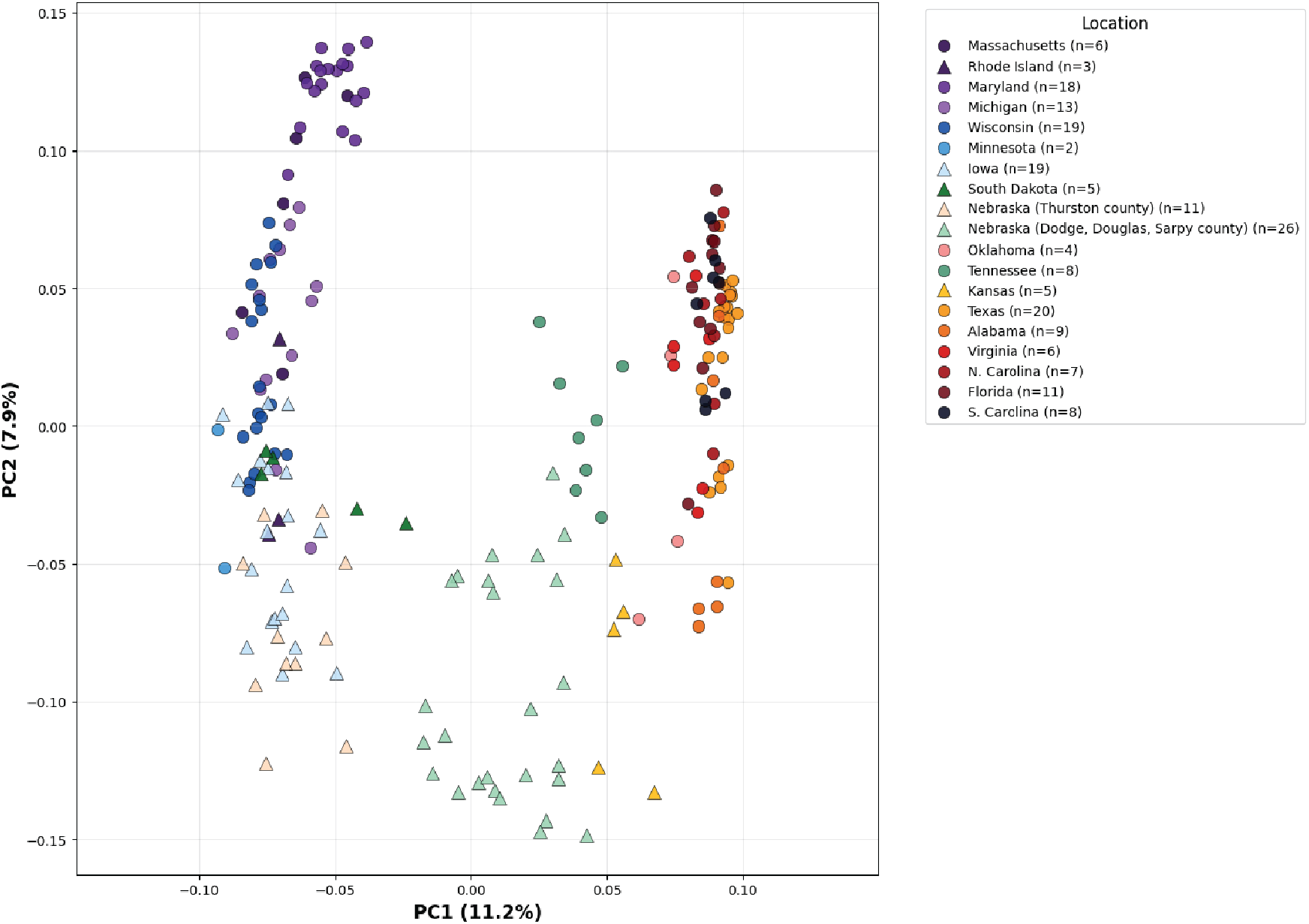
Principal components analysis (PCA) generated with PLINK2 from single nucleotide polymorphism (SNP) datasets for *Ixodes scapularis* based on 20,914 SNPs. Each shape represents a single individual of *I. scapularis*. Shapes are colored by the state of collection for each individual, with Rhode Island and Oklahoma represented by individuals obtained from laboratory colonies rather than field collections. Triangles indicate individuals sequenced as part of this study; circles indicate individuals for which previously-deposited data were downloaded for inclusion.

For DAPC, BIC optimization supported two as the most likely number of genetically distinct clusters, while AIC optimization showed a slight preference for four clusters (**Supplemental Figure 3**). Results from DAPC for both a two-cluster (**Figure 3A**) and four-cluster (**Figure 3B**) analysis are presented here. For the two-cluster analysis, the resulting discriminant function clearly separated specimens from northern versus southern geographic regions with minimal overlap, with most specimens (n=190 of 200; 95%) assigned to one cluster with >94% identity. These cluster assignments largely aligned with the results from PCA, with exceptions for specimens from Tennessee and Nebraska. Specimens from Tennessee grouped confidently with the southern cluster (99.5–100% cluster identity) while specimens from Nebraska showed variation in cluster assignment. All 11 specimens from northeastern Nebraska (Thurston County) and three specimens from central eastern Nebraska (Dodge County) grouped with the northern cluster (97.6–100% cluster identity) and ten specimens from central eastern Nebraska (Dodge, Douglas, or Sarpy Counties) grouped with the southern cluster (94.2– 99.9% identity). The remaining specimens from central eastern Nebraska (n=10) showed mixed cluster assignment, ranging from 85.9% identity with the majority-southern cluster to 89.4% identity with the majority-northern cluster (see **Figure 3A**). In the four-cluster analysis, clusters also aligned with geographic boundaries. Cluster 2 consisted of majority-Northeastern specimens (Rhode Island [BEI], Maryland, and Massachusetts), Cluster 3 consisted of majority-Upper Midwest and northern Great Plains specimens (Michigan, Minnesota, Wisconsin, Iowa, South Dakota, and northeastern Nebraska), and Cluster 4 consisted of majority-southern specimens (Virginia, North Carolina, South Carolina, Alabama, Florida, Texas, and Oklahoma [OSU]) with high cluster assignment identity (>89–99.8%) for nearly all specimens. The exception was a single individual specimen from Massachusetts, which had 73.1% and 26.9% assignment to clusters 3 and 2, respectively. Cluster 1 is composed of specimens from Tennessee, South Dakota, central eastern Nebraska, and Kansas, and a single individual from the OSU laboratory colony, with each individual demonstrating >97.5% cluster assignment identity. This result aligns with the intermediary positioning for specimens from Tennessee, central eastern Nebraska, and Kansas observed in PCA (see **Figure 2**).

**Figure 3.**
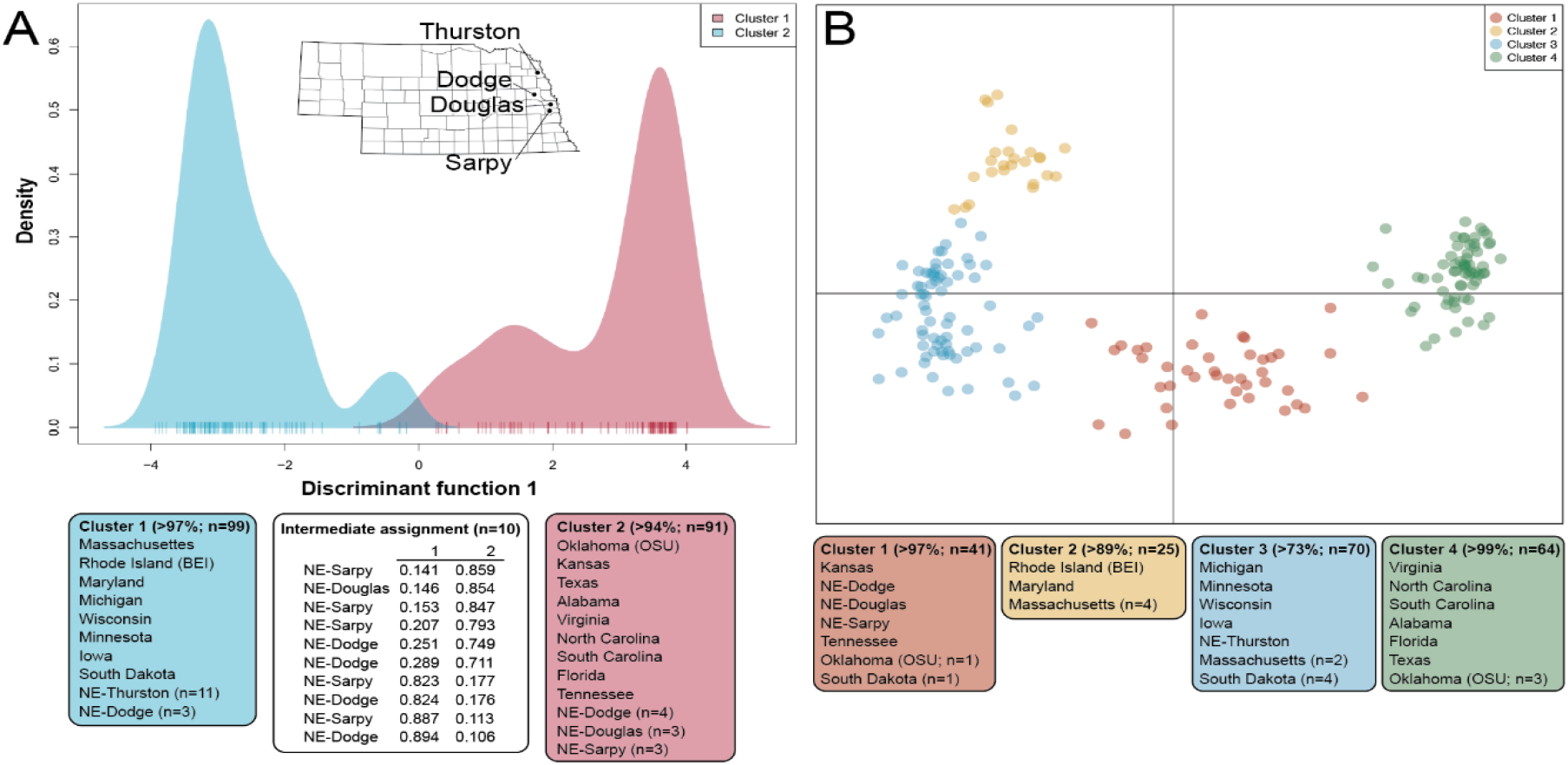
Discriminant analysis of principal components (DAPC) plots from single nucleotide polymorphism (SNP) datasets for *Ixodes scapularis* based on 20,914 SNPs. **(A)** Bayesian information criterion (BlC)-preferred cluster number of two with inset map showing sampling sites within Nebraska labled by county; color-coded lists below plot show individuals assigned to each cluster above a specified threshold of identity; values of n provided for a state indicate that individuals from that population were assinged to multiple clusters; table at center shows individuals from east central Nebraska of intermediate cluster identity, with their county of origin and cluster assignment proportions. **(B)** Akaike information criteria (AlC)-preferred cluster number of four; color-coded lists below plot show individuals assigned to each cluster above a specified threshold of identity; values of n provided for a state indicate that individuals from that population were assinged to multiple clusters.

### Admixture analyses

Analysis of the reduced dataset to compare STRUCTURE and fastSTRUCTURE outputs produced nearly identical results (see **Supplemental Figure 4**), indicating fastSTRUCTURE as a reliable alternative to STRUCTURE for the full dataset. Proportions of ancestry assignment for each individual in the full dataset were largely concordant between fastSTRUCTURE and ADMIXTURE, and aligned with geographic region of origin (**Figure 4**). For fastSTRUCTURE, log-marginal likelihood lower bound (LLBO) estimation supported k=2 while optimal components estimation (KΦ*) supported k=3 (**Supplemental Figure 5**). For ADMIXTURE, cross-validation (CV) errors marginally supported k=3 over k=2 (CV=0.488 versus 0.489, respectively; **Supplemental Figure 5**). Results for both k=2 and k=3 are therefore presented for each analysis (**Figure 4**). For k=2, specimens from the Northeast (Massachusetts, Maryland, Rhode Island [BEI]) and Upper Midwest (Michigan, Wisconsin, Minnesota) shared high proportions of ancestry assignment to a common ancestral population (**Figure 3**; blue or purple bars in k=2 plots). Similarly, specimens from southern states (Texas, Oklahoma [OSU], Alabama, Virginia, North Carolina, South Carolina, and Florida) showed high proportions of ancestry assignment to the second inferred ancestral population (**Figure 3**; red or green bars in k=2 plots). Specimens from Iowa and South Dakota were assigned a majority (>50%) proportion of ancestry to the same ancestral population as Northeast and Upper Midwest specimens, while specimens from Kansas and Tennessee had majority ancestry assignments that aligned with those of specimens from southern states. Specimens from Nebraska displayed clear evidence of admixture, with ancestry assignments varying substantially within the state. Specimens from northeastern Nebraska (Thurston County) had ancestry assignments similar to those of specimens from northern regions, while specimens from central eastern Nebraska (Dodge, Douglas, and Sarpy Counties) displayed mixed ancestry, with assignment proportions near 50% for each individual. Differences in interpretation between k=2 and k=3 were most evident for ADMIXTURE analysis, where addition of a third hypothesized ancestral population revealed support for two ancestral populations contributing to the genomic backgrounds of specimens from northern states, each with varying contributions per individual and roughly distinguishing the Northeast from other populations (**Figure 3**; blue versus yellow bars in k=3 ADMIXTURE plot). For k=3 in the fastSTRUCTURE analysis, addition of a third ancestral population distinguishes specimens from Iowa, South Dakota, Nebraska, and Kansas (**Figure 3**; orange bars in k=3 fastSTRUCTURE plot). For specimens from southern states, interpretation of genetic contribution mostly from a single shared ancestral population was consistent across analyses and values of k.

**Figure 4.**
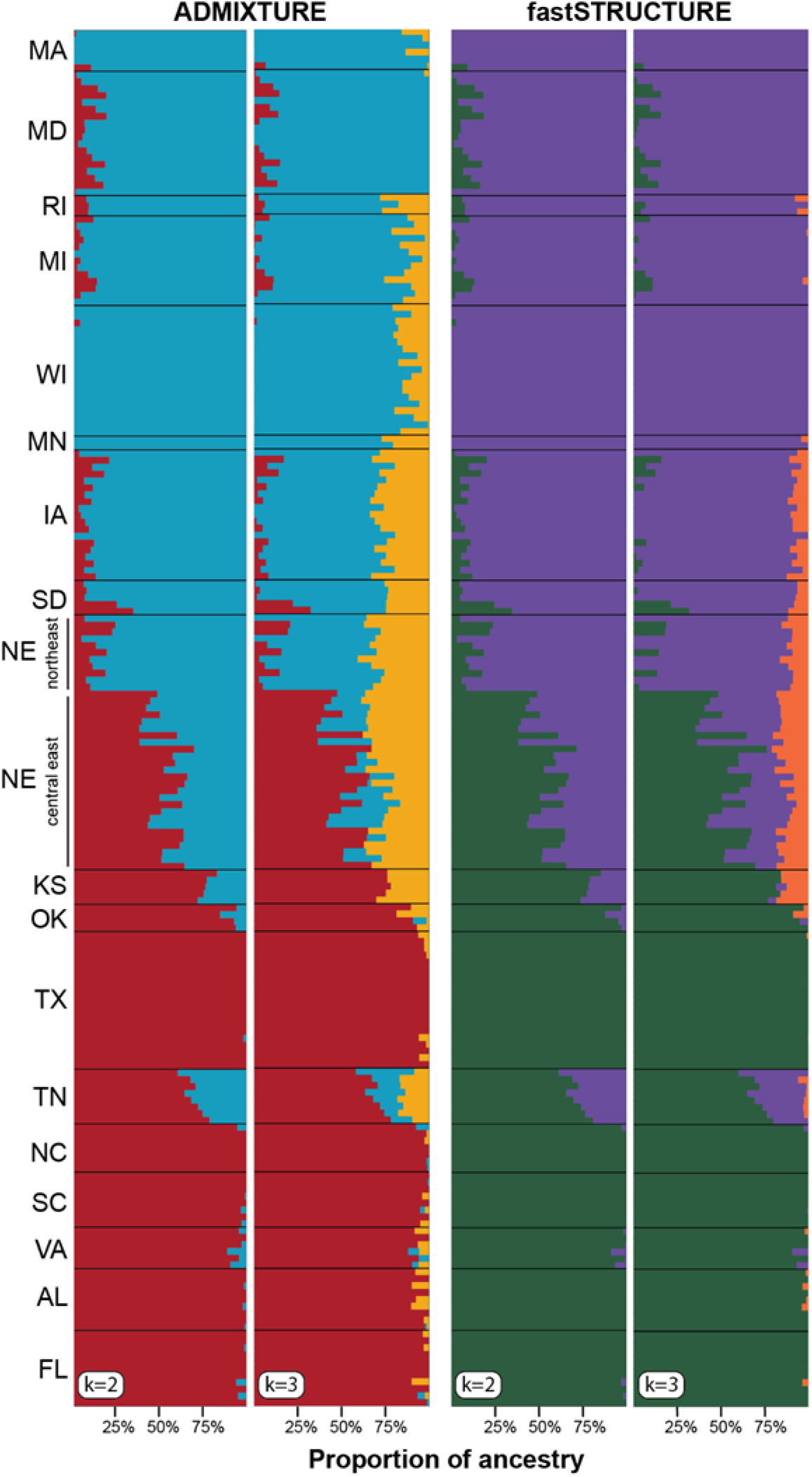
Admixture plots from single nucleotide polymorphism (SNP) datasets for *Ixodes scapularis* based on 20,914 SNPs. Left plots generated using ADMIXTURE; right plots generated using fastSTRUCTURE. For both analyses, the statistically preferred values of k=2 and k=3 for ancestry assignment are shown. Abbreviations for state of origin for individual ticks are given at left

### F_ST_ estimation and visualization of isolation by distance

Corrected F_ST_ values estimated for each pair of populations are summarized in **Table 3**. The greatest F_ST_ values (0.049–0.161; mean 0.1) were estimated between populations with majority-northern ancestry estimation (Rhode Island [BEI], Minnesota, Wisconsin, Michigan, Massachusetts, Maryland, South Dakota, Iowa, and northeastern Nebraska) and majority-southern ancestry estimation (Kansas, Tennessee, Virginia, North Carolina, South Carolina, Alabama, Florida, and Texas). Values among majority-northern ancestry populations (0.002–0.044; mean 0.016) and among majority-southern ancestry populations (0.004–0.046; mean 0.025) were comparatively lower and similar between the two groups, with a higher average F_ST_ estimated among majority-southern ancestry populations. As expected, values were slightly elevated for all comparisons to both the BEI (0.018–0.161; mean 0.074) and OSU (0.007– 0.088; mean 0.049) laboratory colonies. Relatively low F_ST_ values were estimated for all comparisons against the east central Nebraska population (0.014–0.150; mean 0.064), consistent with expectations for genomic admixture in this region. Plotting estimated pairwise F_ST_ values against physical distance between populations to visualize IBD clearly shows elevated F_ST_ values for populations in northern versus southern regions, though R^2^ values (0.1946 and 0.0331, respectively) do not indicate a strong correlation between distance and F_ST_ value for either group compared (**Figure 5**).

**Figure 5.**
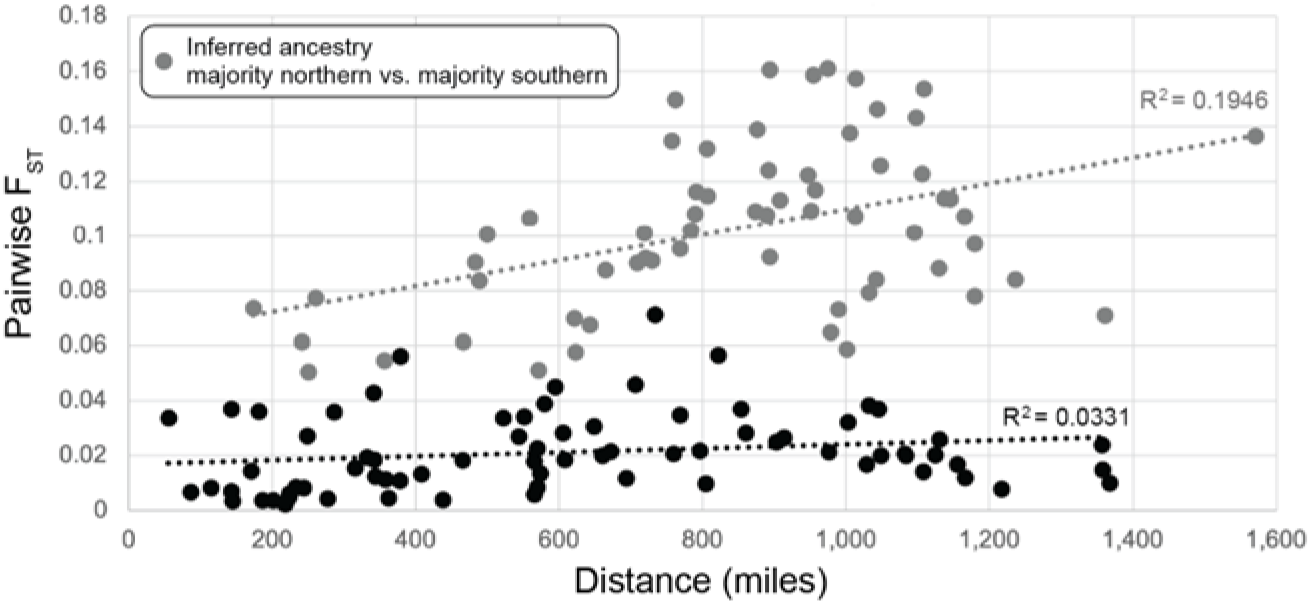
Isolation by distance plot for *Ixodes scapularis*. Pairwise comparisons are made toetween all individuals included in analyses except tor ticks from laboratory-reared colonies (Rhode Island BEI and Oklahoma: OSU). Comparisons between individuals with inferred ancestries that are majority northern vs majority southern according to principal components-anc admixtu^r^e-based analyses are shown in grey. Trendlines for both groups are presented as grey or black dotted lines with R-squared values given for each.

**Table 3.** Corrected pairwise FST values from *Ixodes scapularis* single nucleotide polymorphism (SNP) datasets (n= 92,326 SNPs). Individuals were assigned to populations by state of origin, with individuals from Nebraska desginated as either a northeastern region (NE-NE; Thurston county) or east central region (NE-EC; Dodge. Douglas or Sarpy counties) population. Individuals from laboratory colonies (Rhode Island. BEI: and Oklahoma. OSU) are labeled with their associated laboratory strain acronyms. Greater values are colored a deeper shade of orange.

|  | MN | WI | MI | MA | MD | SD | IA | NE-NE | NE-EC | KS | TN | VA | NC | SC | AL | FL | TX | OK (OSU) |
| --- | --- | --- | --- | --- | --- | --- | --- | --- | --- | --- | --- | --- | --- | --- | --- | --- | --- | --- |
| RI (BEI) | 0.044 | 0.038 | 0.027 | 0.029 | 0.031 | 0.019 | 0.018 | 0.028 | 0.024 | 0.089 | 0.079 | 0.120 | 0.129 | 0.132 | 0.135 | 0.155 | 0.161 | 0.077 |
| MN |  | 0.003 | 0.005 | 0.012 | 0.026 | 0.005 | 0.002 | 0.004 | 0.015 | 0.061 | 0.095 | 0.079 | 0.143 | 0.157 | 0.161 | 0.154 | 0.161 | 0.088 |
| WI |  |  | 0.006 | 0.017 | 0.035 | 0.018 | 0.008 | 0.011 | 0.056 | 0.090 | 0.087 | 0.108 | 0.117 | 0.124 | 0.132 | 0.138 | 0.158 | 0.084 |
| MI |  |  |  | 0.010 | 0.022 | 0.006 | 0.004 | 0.008 | 0.039 | 0.067 | 0.070 | 0.090 | 0.102 | 0.108 | 0.116 | 0.122 | 0.146 | 0.067 |
| MA |  |  |  |  | 0.013 | 0.010 | 0.008 | 0.015 | 0.024 | 0.071 | 0.065 | 0.100 | 0.107 | 0.109 | 0.114 | 0.123 | 0.136 | 0.077 |
| MD |  |  |  |  |  | 0.021 | 0.025 | 0.037 | 0.038 | 0.059 | 0.051 | 0.074 | 0.077 | 0.083 | 0.091 | 0.092 | 0.107 | 0.055 |
| SD |  |  |  |  |  |  | 0.004 | 0.008 | 0.014 | 0.055 | 0.057 | 0.078 | 0.084 | 0.101 | 0.113 | 0.114 | 0.139 | 0.056 |
| IA |  |  |  |  |  |  |  | 0.007 | 0.037 | 0.050 | 0.058 | 0.073 | 0.084 | 0.092 | 0.101 | 0.109 | 0.135 | 0.049 |
| NE-NE |  |  |  |  |  |  |  |  | 0.034 | 0.061 | 0.071 | 0.088 | 0.097 | 0.107 | 0.115 | 0.126 | 0.150 | 0.059 |
| NE-EC |  |  |  |  |  |  |  |  |  | 0.004 | 0.012 | 0.014 | 0.017 | 0.021 | 0.021 | 0.032 | 0.046 | 0.007 |
| KS |  |  |  |  |  |  |  |  |  |  | 0.013 | 0.020 | 0.020 | 0.028 | 0.028 | 0.037 | 0.034 | 0.013 |
| TN |  |  |  |  |  |  |  |  |  |  |  | 0.027 | 0.034 | 0.036 | 0.036 | 0.043 | 0.045 | 0.026 |
| VA |  |  |  |  |  |  |  |  |  |  |  |  | 0.007 | 0.011 | 0.021 | 0.018 | 0.026 | 0.041 |
| NC |  |  |  |  |  |  |  |  |  |  |  |  |  | 0.012 | 0.020 | 0.018 | 0.020 | 0.040 |
| SC |  |  |  |  |  |  |  |  |  |  |  |  |  |  | 0.019 | 0.008 | 0.022 | 0.039 |
| AL |  |  |  |  |  |  |  |  |  |  |  |  |  |  |  | 0.027 | 0.018 | 0.034 |
| FL |  |  |  |  |  |  |  |  |  |  |  |  |  |  |  |  | 0.031 | 0.044 |
| TX |  |  |  |  |  |  |  |  |  |  |  |  |  |  |  |  |  | 0.031 |

### *Borrelia burgdorferi* infection rates inferred from sequencing data

Analysis of *B. burgdorferi* reads in datasets generated for specimens from Iowa, South Dakota, Nebraska, and Kansas revealed heterogeneity in infection rates that correlated with geographic region of origin (**Figure 6**). For ticks from northeastern Nebraska (Thurston County) and Iowa, which consistently grouped with specimens from northern states in PCA- and admixture-based analyses, 8 out of 11 specimens (72.7%) and 8 of 19 specimens (42.1%), respectively, had detectable *B. burgdorferi* reads that aligned across multiple contigs of the reference genome. Specimens from Kansas and South Dakota, despite their different regional affinities in previous analyses (southern versus northern, respectively), showed no detectable *B. burgdorferi* infection (0 out of 5 specimens, each 0%). Ticks from central eastern Nebraska (Douglas, Sarpy, and Dodge Counties), which demonstrate genomic admixture, showed no evidence of *B. burgdorferi* infection (0 out of 29 specimens 0%). All read datasets show some degree of alignment to genomic contig 2 of the *B. burgdorferi* reference genome (Bol26) (Figure 6). Further investigation into specimens with reads aligned to genomic contig 2 only revealed these reads to align to a single ∼900bp region of this contig. Analysis using NCBI BLAST^62^ revealed that this region, and the reads aligning to it across all specimens, shared 100% nucleotide identity to a Rickettsial endosymbiont (GenBank: JQ031633.1) of *I. scapularis* ticks.

**Figure 6.**
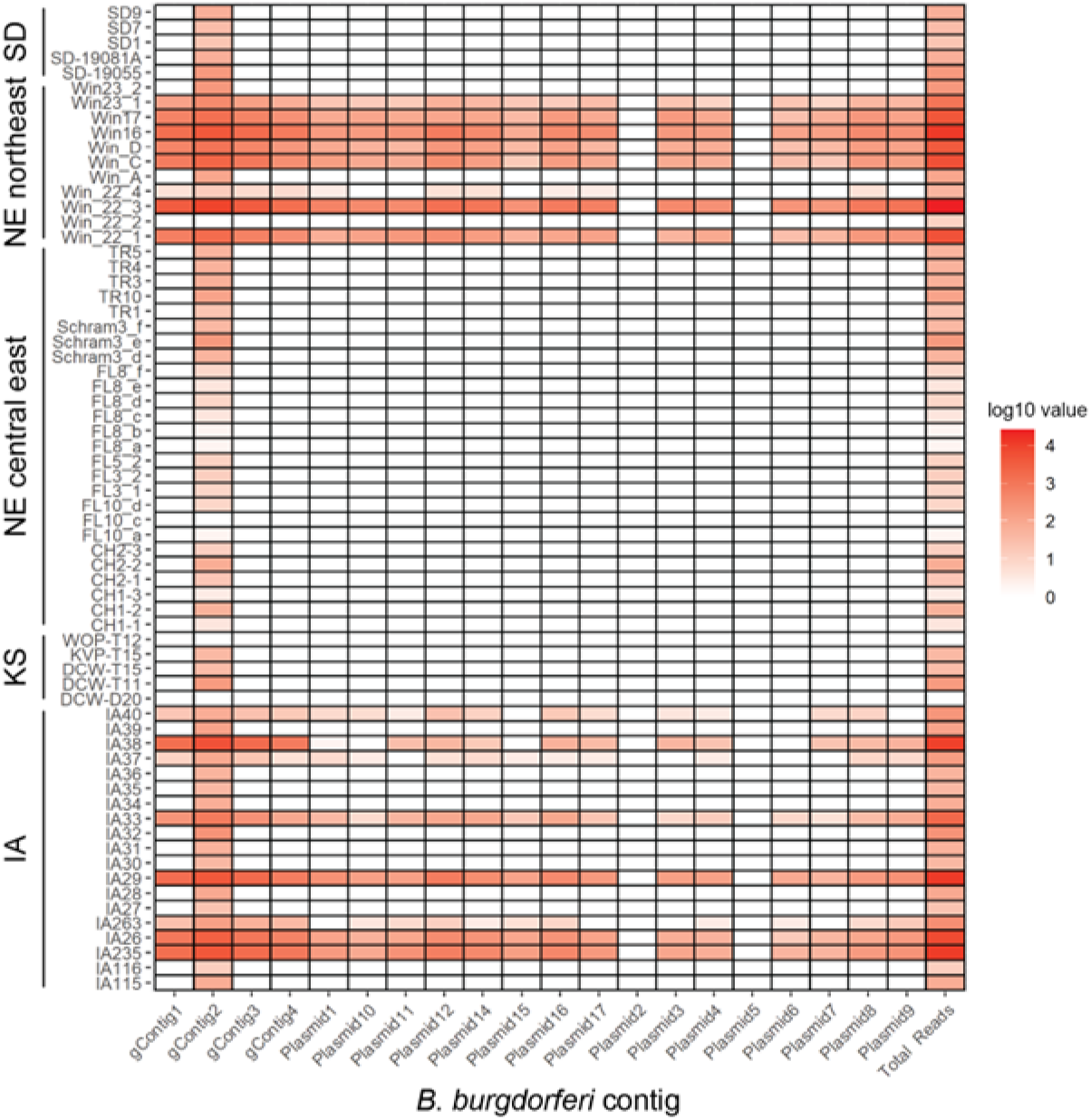
Heatmap of *non-lxodes scapularis* reads that align to contigs of the *Borrelia burgdorferi* reference genome. Read data from individual ticks are shown as rows and are grouped by state of origin; *B. burgdorferi* contigs are shown as columns; plot segements are colored by the number of reads aligned to each contig.

## Discussion

In this study, we leveraged WGS data from individual *I. scapularis* ticks to describe genetic diversity, population structure, admixture, and *B. burgdorferi* infection rates for recently established populations in the Great Plains region of the United States. Dramatically different *B. burgdorferi* infection rates reported for ticks collected in close proximity (<90 miles) within Nebraska^31^ led us to test the hypotheses that 1) Nebraska is a contact zone between historically isolated ancestral populations of *I. scapularis* and 2) strong north-south genetic differentiation and highly structured populations observed in the Eastern United States would also be present in recently-established populations in the Great Plains. Accordingly, we generated 66 novel WGS datasets from individual ticks collected in Nebraska, South Dakota, Iowa and Kansas to compare to publicly available WGS data from ticks collected across much of the Eastern United States. Genomic data show that populations of *I. scapularis* on the western-most edge of the species’ distribution are structured by geography. Admixture occurs in a gradient across the Great Plains, and ticks collected from central eastern Nebraska show the most distinct admixed ancestry, with nearly 50% of the genetic backgrounds of individual ticks being assigned to ancestral northern and southern populations. Presence of *B. burgdorferi* in ticks is easily identified with WGS data and infection rates determined from these data recapitulate local surveillance data.

### Sample selection, sequencing, and variant calling

An objective of this study was to generate and make available WGS data from populations on the western-most edge of the range of *I. scapularis* to provide a more complete picture of the population genomics of this species across the United States (**Figure 1**). Additionally, we aimed to determine the likely origin of recently-established populations of *I. scapularis* in the Great Plains (South Dakota and Nebraska) which have not previously been included in population genomic studies. To put these data into appropriate context, we also generated WGS data from populations of *I. scapularis* in the neighboring states of Iowa and Kansas. While recent studies of *I. scapularis* have included samples from Iowa^15,63^, this study represents the first WGS data from Iowa, South Dakota, Nebraska, and Kansas. These datasets provide more comprehensive sampling of the range of *I. scapularis* by including populations in these zones of recent expansion. Additionally, we generated WGS data for ticks (n=3) from the BEI laboratory colony. These specimens provide additional representation of Northeastern populations for this and future genomic studies, and help to characterize a commonly used laboratory resource. To ensure the accuracy of our results, we employed strict SNP filtration thresholds for the combined dataset of newly-generated and publicly-available data (n=200 specimens). Our final LD-filtered SNP dataset used in PCA- and admixture-based analyses contained ∼21,000 SNPs (see **Table 1**). This falls within the range of the numbers of SNPs analyzed in recent population genomic studies of *I. scapularis* (e.g., from ∼7,000 SNPs in Frederick et al. [2023]^15^ and ∼10,000 SNPs in Dong & Schoville [preprinted]^64^ to <80,000 SNPs in Dong et al. [2025]^63^ and ∼18–67 million in Schoville et al. [2024]^14^). It is worth noting that Schoville et al. imputed missing genotypes in their analysis, which is likely responsible for the greater number of SNPs included in their analyses. The results recovered here for population structure and genomic similarity within and among specimens represented by previously-generated data were largely concordant with those of prior studies, supporting that a sufficient number of loci were sampled here.

### Refined approach for purifying gDNA across sample types

Sample storage and extraction methodology clearly influenced the quantity of gDNA obtained from individual ticks (**Supplemental Figure 2**). Extractions from ticks that were stored frozen following collection consistently produced higher quantities of gDNA compared to ticks of the same life-cycle stage stored in ethanol. This is well-demonstrated by ticks from Iowa, which were frozen following collection, and which produced the highest-quantity gDNA extractions among adult ticks (see **Supplemental Figure 2**). Given the substantially smaller size of nymphal ticks as compared to adults, generating quantities of gDNA sufficient for WGS can be difficult and has previously required strategies such as whole genome amplification^63^. Modifying the Zymo Quick-DNA HWM MagBead Kit protocol allowed us to obtain similar gDNA quantities from nymphal ticks and sets of eight legs removed from adult ticks as compared to whole adults extracted with the Qiagen DNeasy Blood & Tissue Kit protocol. Though extractions from sets of legs were not prepared as sequencing libraries here, the quantities of gDNA obtained from both nymphs and leg sets were sufficient for input to most Illumina library preparation protocols, highlighting the utility of this approach for future studies where input tissue may be limited. Given that the Centers for Disease Control and Prevention (CDC) uses tick bodies for pathogen detection, but does not require legs, our results suggests that tick bodies can be sent to CDC for pathogen testing and their legs retained for generation of genomic data, resulting in reciprocally informative datasets and maximizing the scientific value of each tick collected. This further highlights the flexibility and utility of our modified Zymo Kit-based extraction protocol, which has now been applied successfully to obtain gDNA sufficient for NGS from low-input sample types across multiple species of parasites^34^.

### Population structure and genetic differentiation of *I. scapularis* in the Eastern United States

Aligned with previous studies, our results suggest strong geographic signal for subpopulation structure within *I. scapularis* across the Eastern United States^11–15^. Analyses based on PCA and admixture suggest WGS datasets are best-described by two to four clusters. Division into two clusters, or assuming contribution from two ancestral populations, clearly distinguishes specimens from northern states in the Northeast and Upper Midwest from those from southern states. However, specimens from central eastern Nebraska were not easily distinguished as belonging to one of two clusters at k=2 in the DAPC analysis, and strong genomic admixture was inferred for specimens from both central eastern Nebraska and Tennessee at k=2 in admixture-based analyses (see **Figures 3A; 4**). Schoville et al. also identified evidence of mixed genomic ancestry for specimens from Tennessee, with these specimens similarly falling between two major clusters representing northern and southern populations in their PCA^14^. Results from F_ST_ estimations and visualization of IBD also support genomic distinction between populations in northern and southern states, with slightly elevated F_ST_ values inferred among southern populations, and higher F_ST_ values inferred for north-south pairwise comparisons in nearly all instances (see **Table 3; Figure 5**). Low R-squared values (<0.2) for the plot of F_ST_ versus distance between sampling sites suggest that distance alone does not completely describe this distinction. This further supports the importance of a latitudinal gradient in separation: northern populations at great distances from one another could have relatively low F_ST_ values (e.g., Massachusetts versus Minnesota; ∼1,165 mi apart; F_ST_ 0.01) compared to northern versus southern populations that were closer in physical proximity but demonstrated greater genomic differentiation (e.g., Massachusetts versus Virginia; ∼499 mi apart; F_ST_ 0.1). The range and pattern of pairwise F_ST_ values recovered here was similar compared to previous population genomic studies for *I. scapularis*. Guila-Nuss et al., Frederick et al., and Schoville et al. each found relatively low F_ST_ values among northern populations, slightly elevated values among southern populations, and higher values between the two regions^14,15,65^. This suggests little inbreeding between northern and southern populations, and slightly more isolation and population structure within southern states. At the higher k-values tested here (k=3–4), the Northeast and Upper Midwest (which included northeastern Nebraska and some specimens from South Dakota) were distinguished as separate populations, as was a group variously represented by specimens from more central states, including South Dakota, Iowa, central eastern Nebraska, Kansas and Tennessee (see **Figures 3B; 4**). Previous population genomic studies identified degrees of structure for *I. scapularis*: both Schoville et al. and Dong et al. inferred k=2–3 for their datasets^14,63^, though Guila-Nuss et al. and Frederick et al. each inferred k=5^15,65^. The five clusters of Guila-Nuss et al. were represented by laboratory colony specimens, two southern populations (Florida versus North Carolina and Virginia), and two northern populations (Massachusetts, Maine, and Wisconsin versus New Hampshire and Indiana). The five clusters of Frederick et al. were best-described as Northeastern, Upper Midwestern, Southeastern Atlantic, and Southern Gulf, with central Florida identified as a distinct population. In all, this suggests a gradient of genomic differentiation across the range of *I. scapularis*, with a strong latitudinal signature for subpopulation structure. The two common laboratory strains for *I. scapularis* (BEI, established with ticks from Rhode Island; and OSU, established with ticks from Oklahoma) follow this pattern of latitudinal genomic differentiation, with BEI ticks recovered as more genetically similar to natural northern populations and OSU ticks recovered as more genetically similar to natural southern populations in each of our analyses (see **Figures 2–4**). Therefore, caution is warranted when attempting to draw conclusions about natural populations of *I. scapularis* across the species’ geographic range based on experimental results as these two laboratory populations may also demonstrate differences in behavior and host seeking previously identified in *I. scapularis* across its range, and potentially, as our results suggest, differences in *B. burgdorferi* susceptibility and transmission potential as well (see below).

### Western populations of *I. scapularis* demonstrate a gradient of genomic admixture

By expanding the genomic data available from the western-most edge of the range of *I. scapularis*, we were able to observe admixture occurring at different proportions across a north-south gradient throughout the Great Plains. Under k=2 models, ticks from South Dakota, Nebraska and Kansas displayed evidence of genomic admixture between ancestral northern and southern populations, with the dominant contribution aligning with the latitudinal position of each state. For our DAPC analysis when k=4 (**Figures 3B**), clusters also align with geography, and consist of specimens from Northeastern, Upper Midwestern, and southern states, with Cluster 1 as the exception. This cluster is made up of specimens that demonstrate substantial admixture irrespective of geography, as it includes South Dakota, Nebraska and Kansas, as well as Tennessee (see **Figure 3B**). Patterns of admixture within Nebraska also defy geography. Ticks from northeast Nebraska were inferred to have genomes with predominantly ancestral-northern contributions. Interestingly, ticks from eastern central Nebraska were inferred to have genomes with relatively equal contributions from both northern and southern ancestral populations (see **Figures 3, 4**). This was an unexpected result, as these two Nebraska subpopulations are separated by less than 100 miles, and occur in nearly identical ecoregions. *I. scapularis* has yet to be identified in southeast Nebraska. However, based on the pattern observed here, it seems likely that populations found in this region of the state will have genomic backgrounds that are also admixed, and––like ticks from Kansas––more similar to populations from southern states. As this study represents the first WGS data for ticks from the Great Plains, comparisons to prior studies cannot be made for this region. For Tennessee, which showed admixture proportions comparable to ticks from Kansas and has been included in prior studies, Schoville et al. similarly identified ticks from this state as admixed^14^. However, their demographic analysis suggests that admixture in Tennessee is historical rather than indicative of active gene flow between northern and southern populations. Although we did not conduct a demographic analysis in this study, the identification of genomic admixture in recently-established populations of *I. scapularis* in the Great Plains provides evidence of more recent contact between historically isolated northern and southern populations. Based on analysis of subpopulations within Nebraska, our results also suggest geographic distance alone is not sufficient to describe genomic differentiation within *I. scapularis* across the Eastern United States.

### Identification of *B. burgdorferi* reads from WGS datasets

As expected, for each dataset generated in this study, the vast majority of reads (>98%, **Supplemental Table 1**) aligned to the *I. scapularis* genome. However by aligning the remainder of the reads to the *B. burgdorferi* reference genome, we were able to infer *B. burgdorferi* infection status of *I. scapularis* ticks from WGS datasets alone (**Figure 6**). Presumed infection status largely aligns with public health surveillance data from each state^30,31,66,67^. Samples from Kansas and east central Nebraska did not show evidence of *B. burgdorferi* infection, where >50% of ticks from Iowa and northeastern Nebraska had reads aligned across the *B. burgdorferi* genome. Given that *I. scapularis* ticks from South Dakota have previously been shown to be infected^28,29^, we expected to identify reads aligning to *B. burgdorferi* from these ticks, however our small sample size (N=5) from South Dakota and a lower prevalence of infection from previous studies could explain why reads were not identified. The overall depth of coverage across the *B. burgdorferi* genome was low for any one sample, but the breadth of alignment across the nuclear genome and plasmids increase the reliability of using WGS data to determine infection status. The sequencing effort in this study was aimed at reaching ∼10x depth of coverage, however deeper sequencing of any *B. burgdorferi* positive sample likely would yield more complete genomes suitable for bacterial genomic analysis or phylogenetics. Of note, samples from east central Nebraska that display ∼50% admixture from ancestral northern and southern populations have yet to test positive for *B. burgdorferi* through active surveillance or as a part of this study.

### Study limitations

Sample sizes for several populations included here were relatively low, represented by five or fewer specimens, including Rhode Island (BEI colony; N=3), Oklahoma (OSU colony; N=4), South Dakota (N=5), and Minnesota (N=2). Low sample sizes may have impacted comparative analyses and estimations of summary statistics for these populations. Notably, the Northeast and Lower Midwest regions were also not well-represented in our sampling. Previously-generated datasets available for these regions use reduced representation sequencing approaches^15,65^, and were therefore not included in this study, although a recent analysis provided methodology to blend both WGS and reduced representation datasets^64^. Additionally, we were unable to acquire *I. scapularis* samples from North Dakota where populations have recently established^25^. Given our results suggest genomic admixture in latitudinally-central states like Nebraska and Tennessee, future studies would benefit from inclusion of comparable WGS data from other potential zones of admixture, such as the Ohio River Valley within the Lower Midwest. Finally, it is worth noting that we have assumed all specimens included in analyses are conspecific^68^. Should genomic and phenotypic divergence between populations in northern versus southern regions be deemed sufficient to once again recognize them as separate species^21^, analyses that specifically account for and identify genomic introgression would be a more fitting approach.

## Conclusions

In this study, we expanded the available WGS dataset for *I. scapularis* ticks to include samples from the western-edge of the species’ distribution, resulting in a more representative and complete picture of *I. scapularis* population structure in the Eastern United States. We have shown that there is a strong north-south signal for subpopulation structure in the Great Plains region and that admixture occurs across a gradient. Ticks from central eastern Nebraska demonstrate even admixture from ancestral northern and southern populations and do not show evidence of *B. burgdorferi* infection through WGS datasets or routine surveillance and pathogen testing, consistent with *I. scapularis* populations from the southern U.S. It is imperative to understand how genomic composition affects tick life history traits important for Lyme disease transmission, such as questing behavior, host preference, and vector competence. As *I. scapularis* populations continue to expand and previously isolated populations come into contact, future studies aimed at understanding the consequences of genomic admixture in *I. scapularis* ticks are warranted.

## Acknowledgements

We would like to thank Jordan Rowley, Alok Dhar, Rubayat Khan and others at the UNMC Genomics Core Facility. This work was assisted by UNMC’s Genomics Core Facility which receives partial support from the National Institute of General Medical Sciences (NIGMS) INBRE - P20GM103427 award. This work was supported by Nebraska Research Initiative (NRI) funds under Collaboration Initiative 1114, University of Nebraska System (Project “Nebraska as a Model to Assess Tick-Borne Disease Risk Using Vector Population Genomics,” Application ID 51385) awarded to RC and JRF. This work was partially supported by a grant from the Lyme Disease Association “Pathogen Surveillance and Discovery in Ticks from South Dakota” awarded to JEP. The contents are the sole responsibility of the authors and do not necessarily represent the official views of the NIH or NIGMS. This work was completed utilizing the Holland Computing Center of the University of Nebraska, which receives support from the UNL Office of Research and Innovation, and the Nebraska Research Initiative.

## Literature Cited

1. Centers for Disease Control and Prevention. Surveillance for Ixodes scapularis and pathogens found in this tick species in the United States. https://www.cdc.gov/ticks/resources/TickSurveillance_Iscapularis-P.pdf.

2. Rosenberg, R. et al. Vital Signs: Trends in Reported Vectorborne Disease Cases -United States and Territories, 2004-2016. MMWR Morb. Mortal. Wkly. Rep. 67, 496–501 (2018).

3. CDC. Lyme Disease Surveillance Data. Lyme Disease https://www.cdc.gov/lyme/data-research/facts-stats/surveillance-data-1.html (2025).

4. Hook, S. A. et al. Economic burden of reported Lyme disease in high-incidence areas, United States, 2014-2016. Emerg. Infect. Dis. 28, 1170–1179 (2022).

5. Kugeler, K. J., Schwartz, A. M., Delorey, M. J., Mead, P. S. & Hinckley, A. F. Estimating the frequency of Lyme disease diagnoses, United States, 2010-2018. Emerg. Infect. Dis. 27, 616–619 (2021).

6. CDC. Lyme Disease Case Maps. Lyme Disease https://www.cdc.gov/lyme/data-research/facts-stats/lyme-disease-case-map.html (2025).

7. Eisen, L. & Eisen, R. J. Changes in the geographic distribution of the blacklegged tick, Ixodes scapularis, in the United States. Ticks Tick Borne Dis. 14, 102233 (2023).

8. Humphrey, P. T., Caporale, D. A. & Brisson, D. Uncoordinated phylogeography of Borrelia burgdorferi and its tick vector, Ixodes scapularis. Evolution 64, 2653–2663 (2010).

9. Diuk-Wasser, M. A. et al. Human risk of infection with Borrelia burgdorferi, the Lyme disease agent, in eastern United States. Am. J. Trop. Med. Hyg. 86, 320–327 (2012).

10. Pepin, K. M. et al. Geographic variation in the relationship between human Lyme disease incidence and density of infected host-seeking Ixodes scapularis nymphs in the Eastern United States. Am. J. Trop. Med. Hyg. 86, 1062–1071 (2012).

11. Xu, G., Wielstra, B. & Rich, S. M. Northern and southern blacklegged (deer) ticks are genetically distinct with different histories and Lyme spirochete infection rates. Sci. Rep. 10, 10289 (2020).

12. Sakamoto, J. M., Goddard, J. & Rasgon, J. L. Population and demographic structure of Ixodes scapularis Say in the eastern United States. PLoS One 9, e101389 (2014).

13. Van Zee, J. et al. High SNP density in the blacklegged tick, Ixodes scapularis, the principal vector of Lyme disease spirochetes. Ticks Tick Borne Dis. 4, 63–71 (2013).

14. Schoville, S. D. et al. Genome resequencing reveals population divergence and local adaptation of blacklegged ticks in the United States. Mol. Ecol. 33, e17460 (2024).

15. Frederick, J. C. et al. Phylogeography of the blacklegged tick (Ixodes scapularis) throughout the USA identifies candidate loci for differences in vectorial capacity. Mol. Ecol. 32, 3133–3149 (2023).

16. Ginsberg, H. S. et al. Selective host attachment by Ixodes scapularis (Acari: Ixodidae): Tick-lizard associations in the southeastern United States. J. Med. Entomol. 59, 267–272 (2022).

17. Arsnoe, I., Tsao, J. I. & Hickling, G. J. Nymphal Ixodes scapularis questing behavior explains geographicvariation in Lyme borreliosis risk in the eastern United States. Ticks Tick Borne Dis. 10, 553–563 (2019).

18. Apperson, C. S., Levine, J. F., Evans, T. L., Braswell, A. & Heller, J. Relative utilization of reptiles and rodents as hosts by immature Ixodes scapularis (Acari: Ixodidae) in the coastal plain of North Carolina, USA. Exp. Appl. Acarol. 17, 719–731 (1993).

19. Tietjen, M., Esteve-Gasent, M. D., Li, A. Y. & Medina, R. F. A comparative evaluation of northern and southern Ixodes scapularis questing height and hiding behaviour in the USA. Parasitology 147, 1569–1576 (2020).

20. Durden, L. A., Oliver, J. H., Jr, Banks, C. W. & Vogel, G. N. Parasitism of lizards by immature stages of the blacklegged tick, Ixodes scapularis (Acari, Ixodidae). Exp. Appl. Acarol. 26, 257–266 (2002).

21. Spielman, A., Clifford, C. M., Piesman, J. & Corwin, M. D. Human babesiosis on Nantucket Island, USA: description of the vector, Ixodes (Ixodes) dammini, n. sp. (Acarina: Ixodidae). J. Med. Entomol. 15, 218–234 (1979).

22. Ogden, N. H. et al. Evidence for geographic variation in life-cycle processes affecting phenology of the Lyme disease vector Ixodes scapularis (Acari: Ixodidae) in the United States. J. Med. Entomol. 55, 1386–1401 (2018).

23. Novak, M., Rowley, W., Platt, K., Bartholomew, D. M. & Senne, M. lxodes dammini (Acari, Ixodidae) and Borrelia burgdorferi in Iowa. The Journal of the Iowa Academy of Science: JIAS 98, 99–101 (1991).

24. Coleman, S. The Identification of the Range of Ixodidae Ticks and the Epidemiological Evaluation of Lyme Disease and Spotted Fever Rickettsiosis in Kansas from 2008 to 2012. (Kansas State University, 2013).

25. Russart, N. M., Dougherty, M. W. & Vaughan, J. A. Survey of ticks (Acari: Ixodidae) and tick-borne pathogens in North Dakota. J. Med. Entomol. 51, 1087–1090 (2014).

26. Maestas, L. P., Adams, S. L. & Britten, H. B. First Evidence of an Established Population of Ixodes scapularis (Acari: Ixodidae) in South Dakota. J. Med. Entomol. 53, 965–966 (2016).

27. Nielsen, L. E., Cortinas, R., Fey, P. D., Iwen, P. C. & Nielsen, D. H. First Records of Established Populations of Ixodes scapularis (Acari: Ixodidae) Collected From Three Nebraska Counties. J. Med. Entomol. 57, 939–941 (2020).

28. Black, H., Potts, R., Fiechtner, J., Pietri, J. E. & Britten, H. B. Establishment of Amblyomma americanum populations and new records of Borrelia burgdorferi-infected Ixodes scapularis in South Dakota. J. Vector Ecol. 46, 143–147 (2021).

29. Maestas, L. P., Mays, S. E., Britten, H. B., Auckland, L. D. & Hamer, S. A. Surveillance for Ixodes scapularis (Acari Ixodidae) and Borrelia burgdorferi in Eastern South Dakota State Parks and Nature Areas. J. Med. Entomol. 55, 1549–1554 (2018).

30. Hamik, J. et al. Detection of Borrelia burgdorferi infected Ixodes scapularis (Acari: Ixodidae) associated with local Lyme disease transmission in Nebraska, USA, 2021. Zoonoses Public Health 70, 361–364 (2023).

31. Nebraska Tick Surveillance: Blacklegged Tick. https://experience.arcgis.com/experience/c6fde5fd914d407aad18c07cc7725d6a/page/Blacklegged-Tick?views=Lyme-Disease#data_s=id%3AdataSource_14-18c922b9e53-layer-3%3A2%2Cid%3AdataSource_9-18c922b9e53-layer-3%3A64.

32. Ng’eno, E. et al. Phenology of five tick species in the central Great Plains. PLoS One 19, e0302689 (2024).

33. McClung, K. L., Sundstrom, K. D., Lineberry, M. W., Grant, A. N. & Little, S. E. Seasonality of Amblyomma americanum (Acari: Ixodidae) activity and prevalence of infection with tick-borne disease agents in north central Oklahoma. Vector Borne Zoonotic Dis. 23, 561–567 (2023).

34. Herzog, K. S., Wu, R., Hawdon, J. M., Nejsum, P. & Fauver, J. R. Assessing de novo parasite genomes assembled using only Oxford Nanopore Technologies MinION data. iScience 27, 110614 (2024).

35. National Center for Biotechnology Information. NCBI SRA Toolkit. (Github).

36. Chen, S., Zhou, Y., Chen, Y. & Gu, J. fastp: an ultra-fast all-in-one FASTQ preprocessor. Bioinformatics 34, i884–i890 (2018).

37. Bioinformatics, B. FastQC A Quality Control tool for High Throughput Sequence Data. https://www.bioinformatics.babraham.ac.uk/projects/fastqc/.

38. De, S. et al. A high-quality Ixodes scapularis genome advances tick science. Nat. Genet. 55, 301–311 (2023).

39. RepeatMasker. (Github).

40. Li, H. & Durbin, R. Fast and accurate short read alignment with Burrows-Wheeler transform. Bioinformatics 25, 1754–1760 (2009).

41. Li, H. et al. The Sequence Alignment/Map format and SAMtools. Bioinformatics 25, 2078–2079 (2009).

42. Picard. https://broadinstitute.github.io/picard/.

43. McKenna, A. et al. The Genome Analysis Toolkit: a MapReduce framework for analyzing next-generation DNA sequencing data. Genome Res. 20, 1297–1303 (2010).

44. DePristo, M. A. et al. A framework for variation discovery and genotyping using next-generation DNA sequencing data. Nat. Genet. 43, 491–498 (2011).

45. Van der Auwera, G. A. et al. From FastQ data to high confidence variant calls: the Genome Analysis Toolkit best practices pipeline. Curr. Protoc. Bioinformatics 43, 11.10.1–11.10.33 (2013).

46. Danecek, P. et al. The variant call format and VCFtools. Bioinformatics 27, 2156–2158 (2011).

47. Catchen, J., Hohenlohe, P. A., Bassham, S., Amores, A. & Cresko, W. A. Stacks: an analysis tool set for population genomics. Mol. Ecol. 22, 3124–3140 (2013).

48. Rochette, N. C., Rivera-Colón, A. G. & Catchen, J. M. Stacks 2: Analytical methods for paired-end sequencing improve RADseq-based population genomics. Mol. Ecol. 28, 4737–4754 (2019).

49. Chang, C. C. et al. Second-generation PLINK: rising to the challenge of larger and richer datasets. Gigascience 4, 7 (2015).

50. Hunter, J. D. Matplotlib: A 2D Graphics Environment. Comput. Sci. Eng. 9, 90–95 (2007).

51. Waskom, M. seaborn: statistical data visualization. J. Open Source Softw. 6, 3021 (2021).

52. Jombart, T. adegenet: a R package for the multivariate analysis of genetic markers. Bioinformatics 24, 1403–1405 (2008).

53. Jombart, T. & Ahmed, I. adegenet 1. 3-1: new tools for the analysis of genome-wide SNP data. Bioinformatics 27, 3070–3071 (2011).

54. R Core Team. R: A language and environment for statistical computing. R Foundation for Statistical Computing, Vienna, Austria. http://www.R-project.org/.

55. Raj, A., Stephens, M. & Pritchard, J. K. fastSTRUCTURE: variational inference of population structure in large SNP data sets. Genetics 197, 573–589 (2014).

56. Alexander, D. H., Novembre, J. & Lange, K. Fast model-based estimation of ancestry in unrelated individuals. Genome Res. 19, 1655–1664 (2009).

57. Pritchard, J. K., Stephens, M. & Donnelly, P. Inference of population structure using multilocus genotype data. Genetics 155, 945–959 (2000).

58. Falush, D., Stephens, M. & Pritchard, J. K. Inference of population structure using multilocus genotype data: linked loci and correlated allele frequencies. Genetics 164, 1567–1587 (2003).

59. Lischer, H. E. L. & Excoffier, L. PGDSpider: an automated data conversion tool for connecting population genetics and genomics programs. Bioinformatics 28, 298–299 (2012).

60. Evanno, G., Regnaut, S. & Goudet, J. Detecting the number of clusters of individuals using the software STRUCTURE: a simulation study. Mol. Ecol. 14, 2611–2620 (2005).

61. Earl, D. A. & vonHoldt, B. M. STRUCTURE HARVESTER: a website and program for visualizing STRUCTURE output and implementing the Evanno method. Conserv. Genet. Resour. 4, 359–361 (2012).

62. Altschul, S. F., Gish, W., Miller, W., Myers, E. W. & Lipman, D. J. Basic local alignment search tool. J. Mol. Biol. 215, 403–410 (1990).

63. Dong, D.-Y., Paskewitz, S. M., Tsao, J. I. & Schoville, S. D. Genetic and landscape connectivity of Blacklegged ticks during range expansion in select states of the Midwestern USA. Ecol. Evol. 15, e72360.(2025).

64. Dong, D.-Y. & Schoville, S. D. Reconstructing the demographic history of blacklegged ticks (Ixodes scapularis) in the northern United States. bioRxiv 2026.03.10.710853 (2026) doi:10.64898/2026.03.10.710853.

65. Gulia-Nuss, M. et al. Genomic insights into the Ixodes scapularis tick vector of Lyme disease. Nat. Commun. 7, 10507 (2016).

66. Lingren, M., Rowley, W. A., Thompson, C. & Gilchrist, M. Geographic distribution of ticks (Acari: Ixodidae) in Iowa with emphasis on Ixodes scapularis and their infection with Borrelia burgdorferi. Vector Borne Zoonotic Dis. 5, 219–226 (2005).

67. Oliver, J. D., Bennett, S. W., Beati, L. & Bartholomay, L. C. Range Expansion and Increasing Borrelia burgdorferi Infection of the Tick Ixodes scapularis (Acari: Ixodidae) in Iowa, 1990-2013. J. Med. Entomol. 54, 1727–1734 (2017).

68. Oliver, J. H., Jr et al. Conspecificity of the ticks Ixodes scapularis and I. dammini (Acari: Ixodidae). J. Med. Entomol. 30, 54–63 (1993).

