## Supplemental Figures for "When clades collide: Genomic admixture in blacklegged ticks (*Ixodes scapularis*) from the Great Plains"

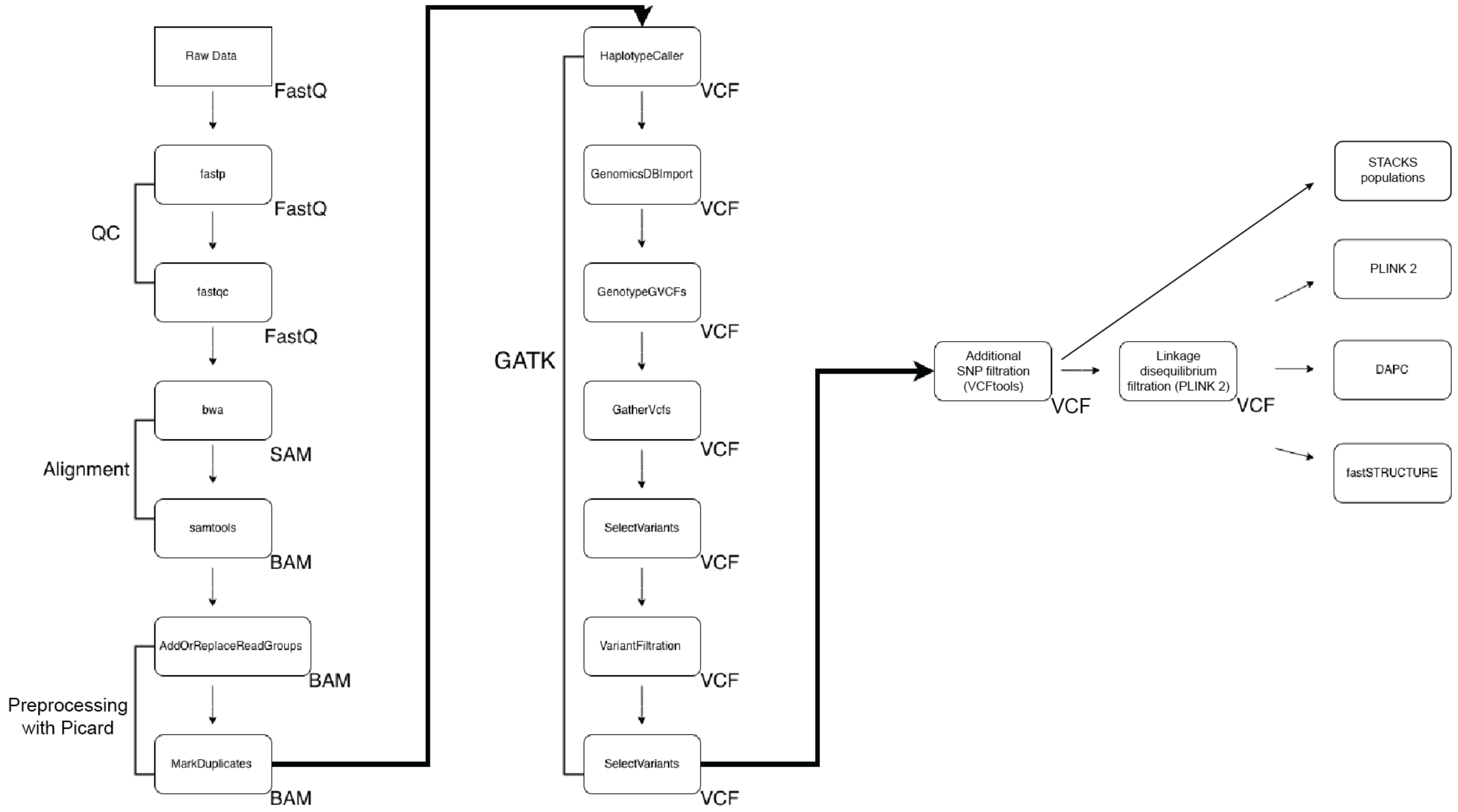


**Supplemental Figure 1. Schematic of bioinformatic pipeline used to generate, filter, and analyze single nucleotide polymorphism (SNP) datasets for individuals of *Ixodes scapularis*.**

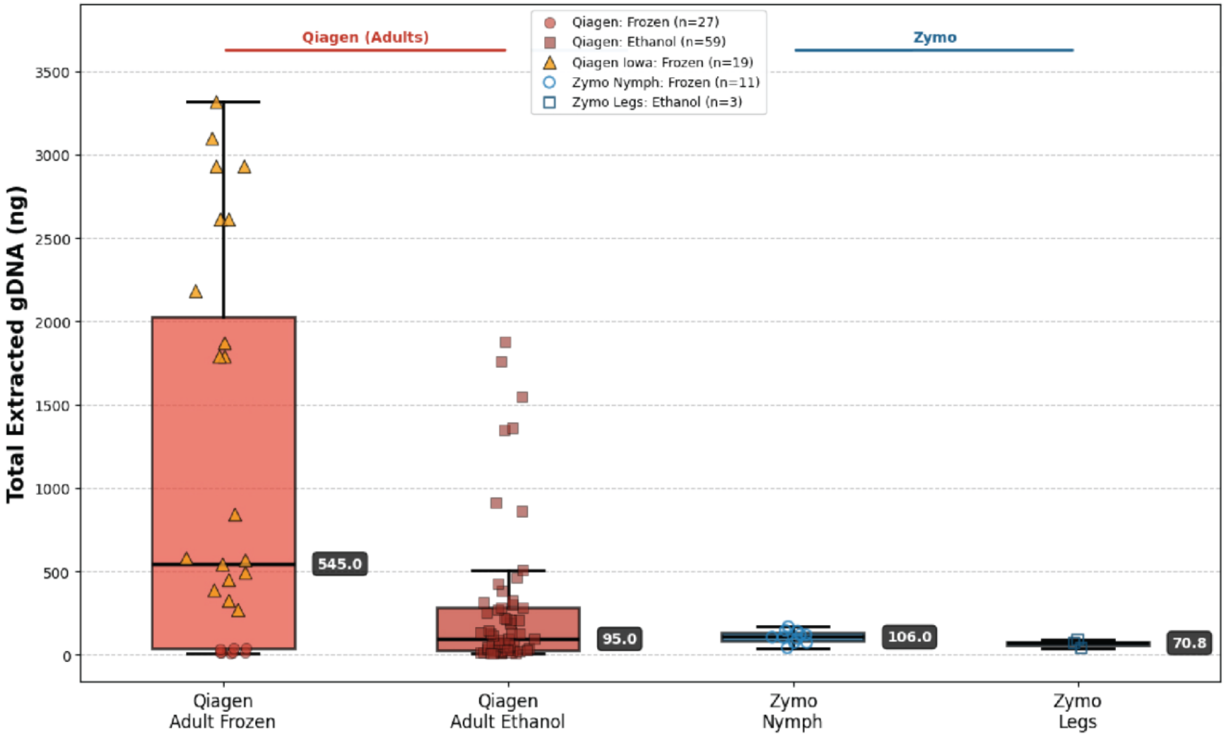


**Supplemental Figure 2. Summary of DNA extraction efficiency from individual *Ixodes scapularis* sequenced as part of this study compared by amount of input tissue, preservation method, and extraction method.** The average mass of DNA purified from each individual using each method (Qiagen DNEasy Blood & Tissue Kit vs. Zymo Quick-DNA HWM MagBead Kit) is provided in dark gray to the right of each box-and-whisker plot. The modified Zymo protocol performed comparably to the modifed Qiagen protocol in terms of total average mass of DNA purified from an individual tick, especially considering the Zymo protocol was only used with nymphal ticks or the legs of ticks from a single adult while the Qiagen protocol was only used with whole adult ticks. Adult ticks collected from Iowa were outliers in terms of total mass of DNA purified from each individual.

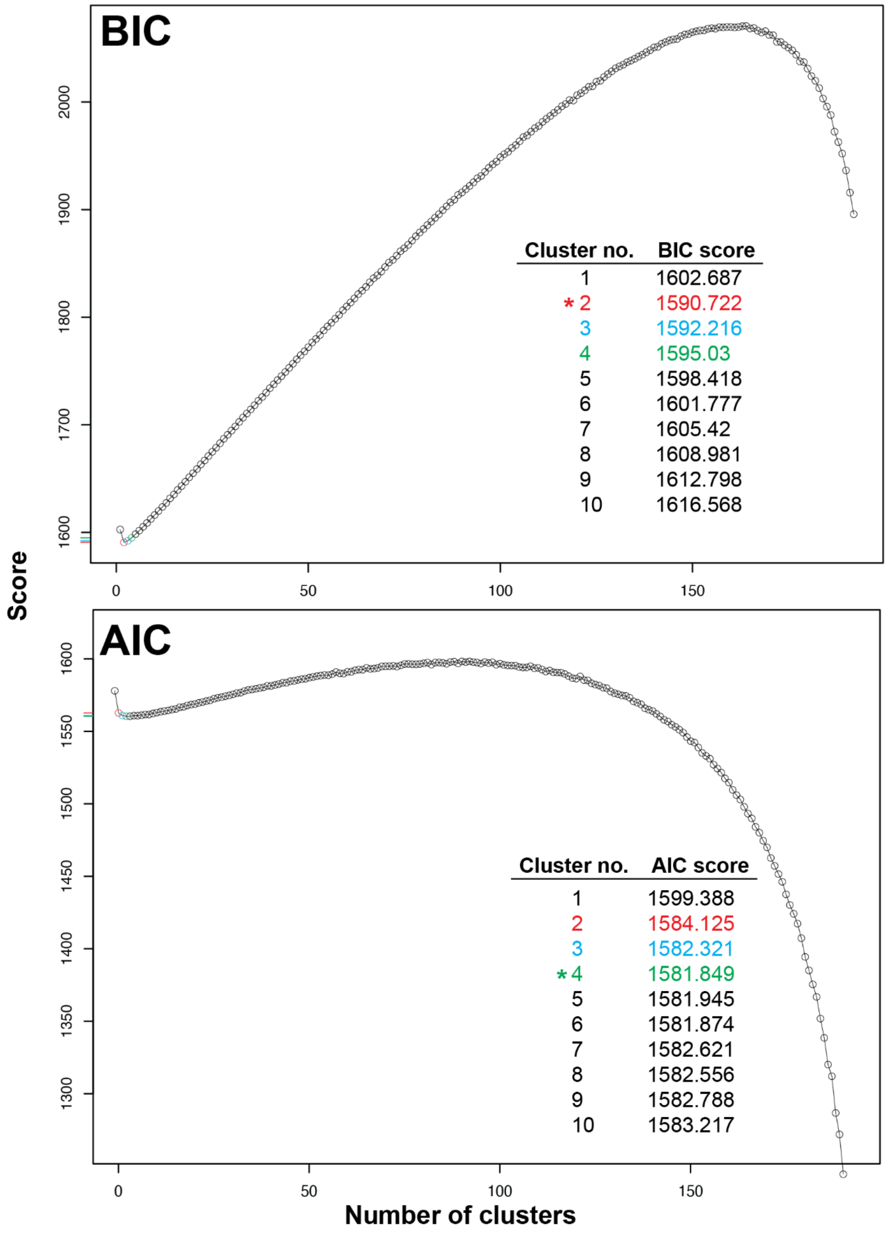


**Supplemental Figure 3. Bayesian information criterion (BIC) and Akaike information criterion (AIC) scores versus cluster number for DAPC analysis.** Scores for cluster numbers of k=2 (red), k=3 (blue), and k=4 (green) are highlighted on each graph. Lowest score for each criterion is noted with asterisk.

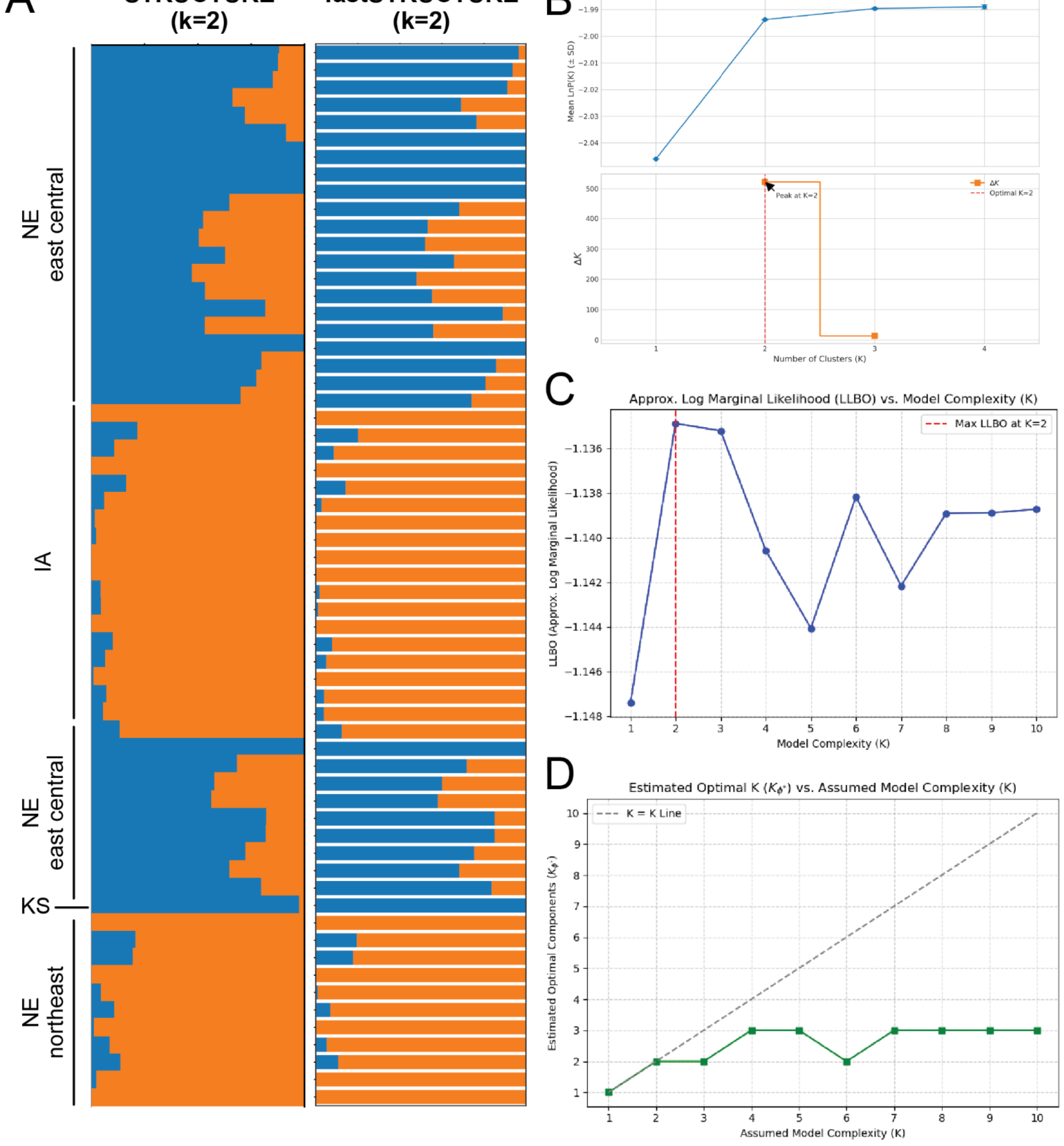


**Supplemental Figure 4. Demonstration of congruence between STRUCTURE and fastSTRUCTURE results for analysis of a reduced single nucleotide polymorphism (SNP) dataset representing a subset of individuals of *Ixodes scapularis* from teh Great Plains newly-sequenced as part of this study. (A)** Admixture plots for STRUCTURE (left) and fastSTRUCTURE (right); **(B)** STRUCTURE Harvester k-value [illegible]

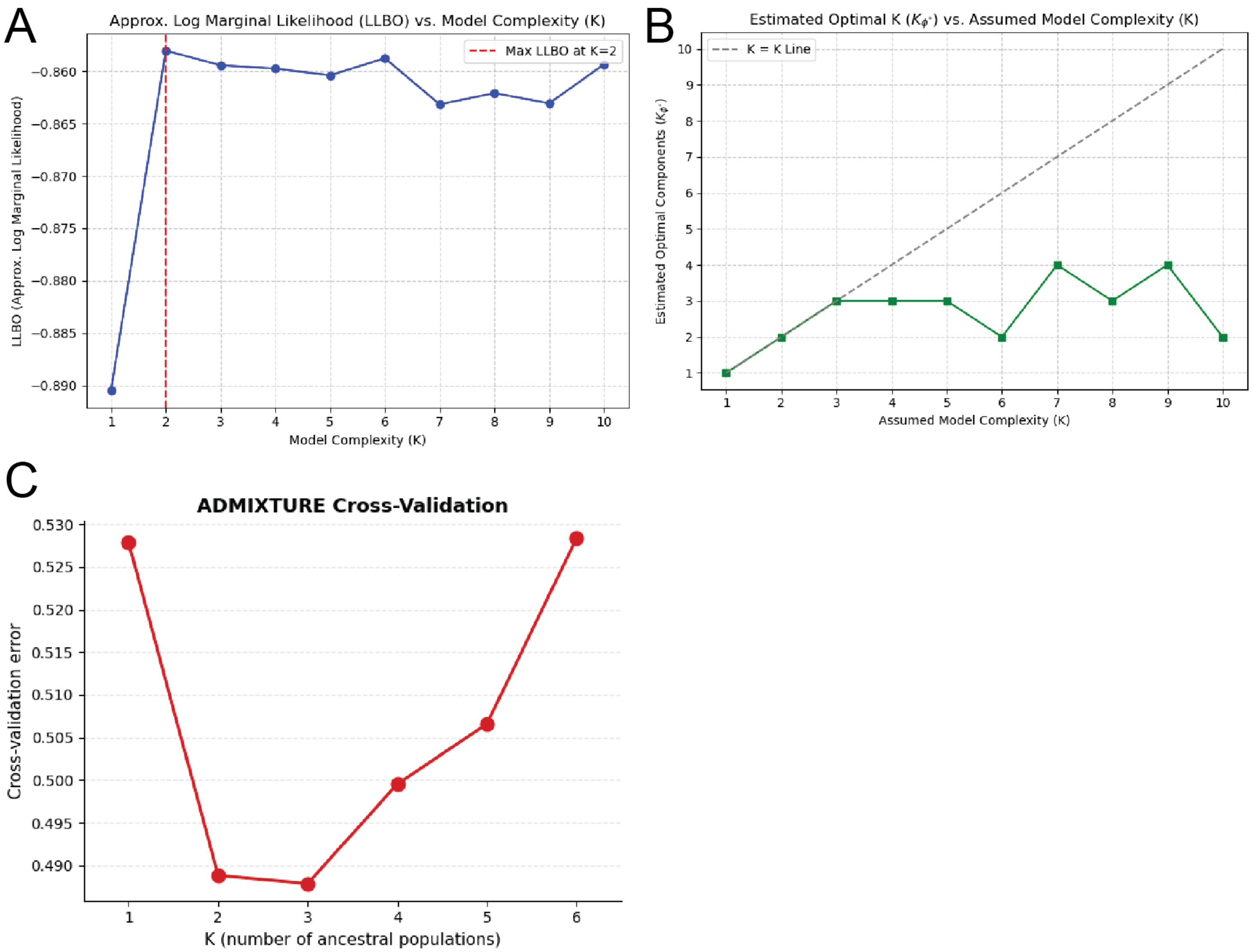


**Supplemental Figure 5. k-value estimation results for fastSTRUCTURE (A, B) and ADMIXTURE (C).** Based on the complete *Ixodes scapularis* single nucleotide polymorphism (SNP) dataset (20,914 SNPs; n=200 individuals).
